# Structural basis for tetherin antagonism as a barrier to zoonotic lentiviral transmission

**DOI:** 10.1101/638890

**Authors:** Cosmo Z. Buffalo, Christina M. Stürzel, Elena Heusinger, Dorota Kmiec, Frank Kirchhoff, James H. Hurley, Xuefeng Ren

## Abstract

Tetherin is a host defense that physically prevents escape of virions from the plasma membrane. Human tetherin lacks the motif DIWK antagonized by SIV, the antecedent of HIV. Here, we reconstituted the AP-2 clathrin adaptor complex with a simian tetherin and SIV Nef and determined its structure by cryo-EM. Nef refolds the first α-helix of the β2 subunit of AP-2 to a β hairpin, creating a binding site for the DIWK sequence. The tetherin binding site in Nef is distinct from those of MHC-I, CD3, and CD4, but overlaps the site for SERINC5 restricting viral infectivity. The structure explains the dependence of SIVs on the host tetherin DIWK sequence and the consequent barrier to human transmission.

## Introduction

Human and simian immunodeficiency viruses (HIV and SIV) depend on factors known as accessory viral proteins for their efficient replication and spread (*1*). Lentiviral accessory proteins, including Nef, Vpu, Vif, Vpr, and Vpx, counteract host defenses by redirecting cellular pathways for the benefit of the virus (*1, 2*). In some cases, activities of these viral proteins overlap or compensate for one another to ensure viral fitness. The counteraction of the host factor tetherin by lentiviral Nef, Env, and Vpu is a prominent example (*3*). Tetherin (also known as BST2) is an interferon-induced antiviral restriction factor that physically tethers nascent virions to the plasma membrane of the viral producer cell, preventing release and marking the cell for immune detection (*4*). The sequences of tetherin orthologs vary considerably between even closely related species and tetherin is counteracted by viral accessory proteins in a species-specific manner (*5, 6*). Most notably, the lack of an otherwise conserved (G/D)DIWK motif in human tetherin confers resistance to downregulation by the viral protein Nef used by most SIVs to counteract this restriction factor. Thus, antagonism of tetherin is a major barrier to successful simian-human transmission of non-human primate lentiviruses, the origin of HIV (*7*). Here, we elucidate the structural basis for antagonism of (G/D)DIWK-containing simian tetherin by SIV Nef in near-atomic detail.

The accessory proteins Nef and Vpu of HIV and SIV reconfigure the membrane protein repertoire of host cells by hijacking clathrin-coated vesicles (CCVs) to sequester and/or degrade target cargoes (*8*). The heterotetrameric adaptor protein (AP) complexes are primarily responsible for connecting cargo and membranes to clathrin. The trans-Golgi network (TGN) and plasma membrane complexes AP-1 and AP-2, respectively, are major direct targets of Nef and Vpu (*1, 8*). APs consists of two large solenoidal subunits (α or γ and β), a medium subunit μ, and a small subunit σ (*9*). All known host cargoes interact with APs via either tyrosine-based sorting signals (YxxF) that bind to the C-terminal domain (CTD) of μ or dileucine signals (ExxL(L/V)) that bind to a hemicomplex of α/γ and σ (*9*). Nef and Vpu both use (ExxxL(L/V)) motifs to hijack AP complexes (*10–13*). In solution, these sites are hidden in the locked conformation of APs (*14, 15*). Upon activation by the small GTPase Arf1 at TGN membranes (*16*) or the lipid PI(4,5)P_2_ at the plasma membrane (PM) (*14*), the μ CTD undergoes a large conformational change, unlocking and activating the complex.

To overcome barriers to cross-species transmission during primate lentiviral evolution, the task of antagonizing tetherin has shifted repeatedly between Nef and Vpu (*7*). Most SIV including SIVcpz, SIVgor, and SIVsmm, the direct precursors of HIV-1 and HIV-2, respectively, use their Nef protein to target AP-2 at the PM by a mechanism whose details are still unknown (*3, 6, 17, 18*). Pandemic group M HIV-1 strains switched from Nef to Vpu to hijack AP-1 at the TGN, using its dileucine motif to unlock AP-1 and free the μ CTD to target a variant Tyr motif in the cytosolic tail of tetherin (*13, 19*). In contrast, Nef proteins from the epidemic HIV-1 group O evolved to interact with a region in human tetherin sequence adjacent to the (G/D)DIWK deletion bridging the μ and σ subunits of AP-1 in *trans* and promoting a unique trimeric assembly of AP-1 to sequester tetherin at the TGN (*20, 21*). The remaining N and P groups of HIV-1 have gained little if any activity against human tetherin and were only detected in a few individuals. Here, we report on the Nef of SIVsmm infecting sooty mangabeys. SIVsmm uses Nef and its envelope (Env) glycoprotein as tetherin antagonist in is natural simian host and represents the direct ancestor of HIV-2 and of SIVmac causing simian AIDS in rhesus macaques (*7*). SIVsmm managed to cross the species barrier from monkeys to humans on at least 9 occasions possibly because its Env protein shows some activity against human tetherin (*22*). However, the Nef proteins of HIV-2 gained little if any activity against human tetherin (*22, 23*) and HIV-2 shows lower transmission rates and is much less prevalent than HIV-1. We show by cryo-EM and allied methods that simian tetherin is targeted in a unique way by the induced refolding of the β subunit of AP-2 from an α-helix to a β-hairpin, creating a novel binding site for the (G/D)DIWK motif in a crevice between the new β-hairpin and Nef.

## Results

### Cooperative interaction between AP-2, simian tetherin and SIVsmm Nef

In the absence of the lipid membrane, the AP-2 core adopts a locked conformation that obscures the binding sites for recruiting both physiological cargoes and Nef (Fig. 1A). In order to promote the open conformation of AP-2 in the absence of lipid membranes, we generated the unlocked tetrameric core construct AP2^ΔµC^, in which the µ2 CTD that is dispensable for Nef binding to AP-2 (*24*), is deleted. We compared the SIVsmm Nef and simian tetherin binding of this construct to that of the AP-2 α-σ2 hemicomplex (“AP2^hemi^“). AP2^hemi^ binds tightly to HIV-1 NL4-3 Nef in the absence of cargo (*25*), but binds only very weakly to SIVmac239 Nef (96-237) and rhesus (rh) tetherin (*26*). By tetherin tail-MBP pull down assay, we observed a tight cooperative interaction between rh tetherin, SIVsmm Nef and AP2^ΔµC^ in combination. In contrast, the tetherin tail did neither bind to AP2^ΔµC^ or SIVsmm Nef alone nor interact with AP2^hemi^, even in the presence of SIVsmm Nef (Fig 1B). We used fluorescence polarization (FP) to determine that Alexa488-tetherin tail bound to AP2^ΔµC^ and SIVsmm Nef with *K*d = 2.7 µM ± 0.1 µM (Fig 1C, 1D), while binding to the separate components or to AP2^hemi^ was undetectable. These data indicate that simian tetherin binds to the AP-2 core and SIVsmm Nef in a cooperative and high affinity ternary complex that requires elements of the full AP-2 tetramer.

**Fig. 1.**
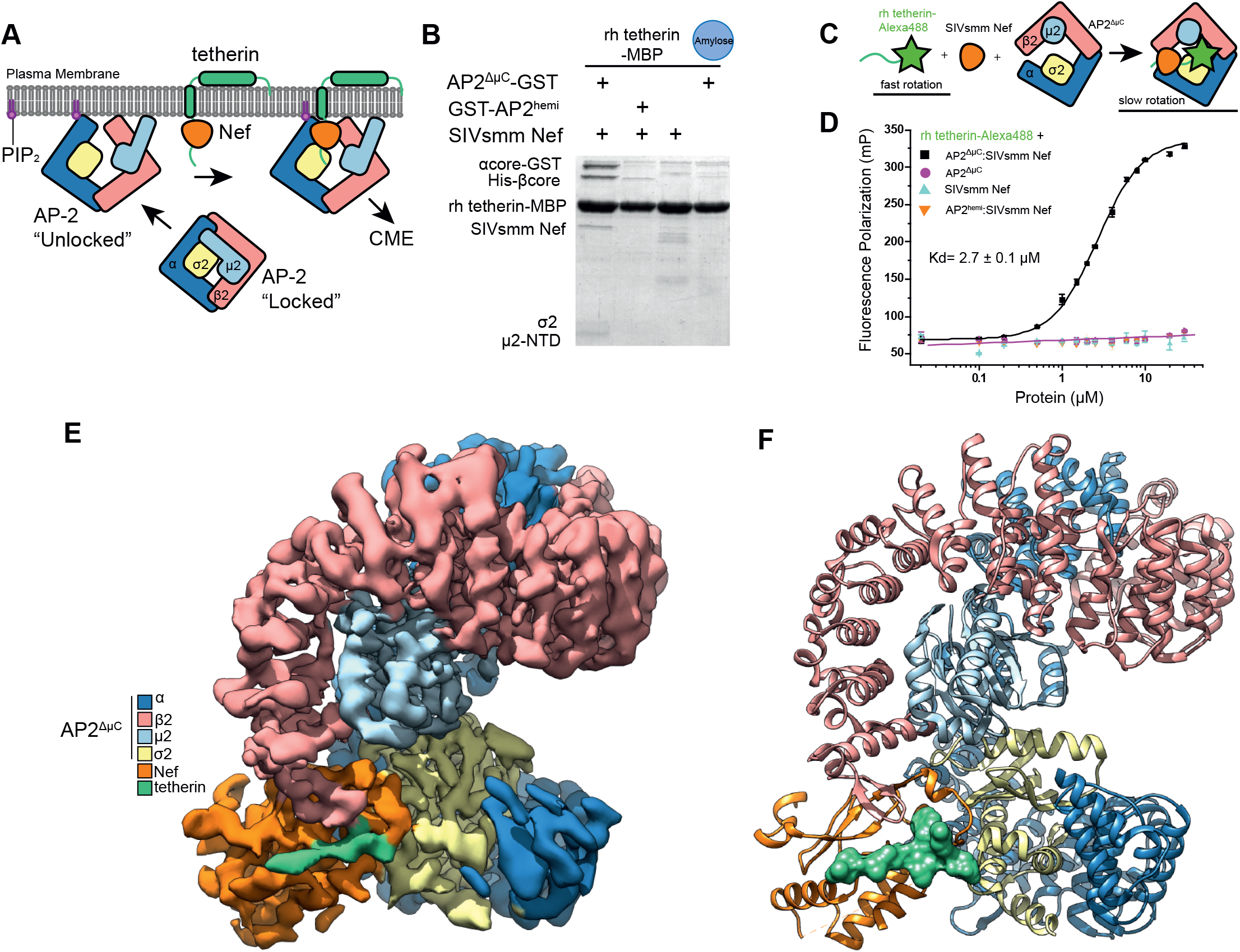
Reconstitution and atomic model of the AP2^ΔµC^:smm tetherin-SIVsmm Nef complex. **(A)** Cartoon representation of AP-2 unlocking at the plasma membrane when it interacts with PIP^2^. Nef can now recruit tetherin to the unlocked AP-2 complex where clathrin mediated endocytosis (CME) will result in Nef-induced downregulation of tetherin from the plasma membrane. (**B**) Simian tetherin tail synergistically binds to AP-2 and SIVsmm Nef. Rh tetherin-MBP was immobilized in amylose resin to pull down SIVsmm Nef and/or AP2^ΔµC^, α-σ2 hemi complex. **(C)** Upper panel: Cartoon representation of FP assay. Alexa488 labeled tetherin tail rotates much slower and further causes FP signal increasing while binding to large AP2^ΔµC^ and SIVsmm Nef. Lower panel: Fluorescence polarization result indicates the cooperative interaction between AP2^ΔµC^, rh tetherin and SIVsmm Nef. Fluorescence polarization (in units of mP) is plotted as a function of AP2^ΔµC^ /SIVsmm Nef concentration (µM) using a logarithmic scale. **(D)** Cryo-EM density map of AP2^ΔµC^ bound to SIVsmm Nef and the smm tetherin cytoplasmic tail **(E)** Ribbon model of AP2^ΔµC^ bound to SIVsmm Nef and the smm tetherin cytoplasmic tail build from the AP2^ΔµC^:smm tetherin-SIVsmm Nef cryo-EM density map.

### CryoEM structure of the AP2^ΔµC^, smm tetherin and SIVsmm Nef complex

The structure of the complex of AP2^ΔµC^ cross-linked to the tetherin-SIVsmm Nef complex was determined by cryo-EM at a resolution of 3.8 Å (Fig. 1E, F, S1, S2). Side-chain density was visible throughout most of the density map, including most of all four subunits of AP-2, SIVsmm Nef, and smm tetherin. An atomic model was built on the basis of the crystal structures of unlocked AP-2 (*14*) and NL4-3 HIV-1 Nef bound to AP2^hemi^ (*25*). The tetherin tail and unique regions of SIVsmm Nef were built *ab initio* (Fig. 2A-C). The conformation of the dileucine loop in the cross-linked SIVsmm Nef complex is essentially identical to the conformation previously seen in the uncrosslinked NL4-3 complex (*25*) (Fig. 2D), showing that the presence of the crosslink did not perturb the structure. The overall conformation of AP2^ΔµC^ is similar to that of the unlocked AP-2 core (*14*), with the pivotal exception of the β2 N-terminus.

**Fig. 2.**
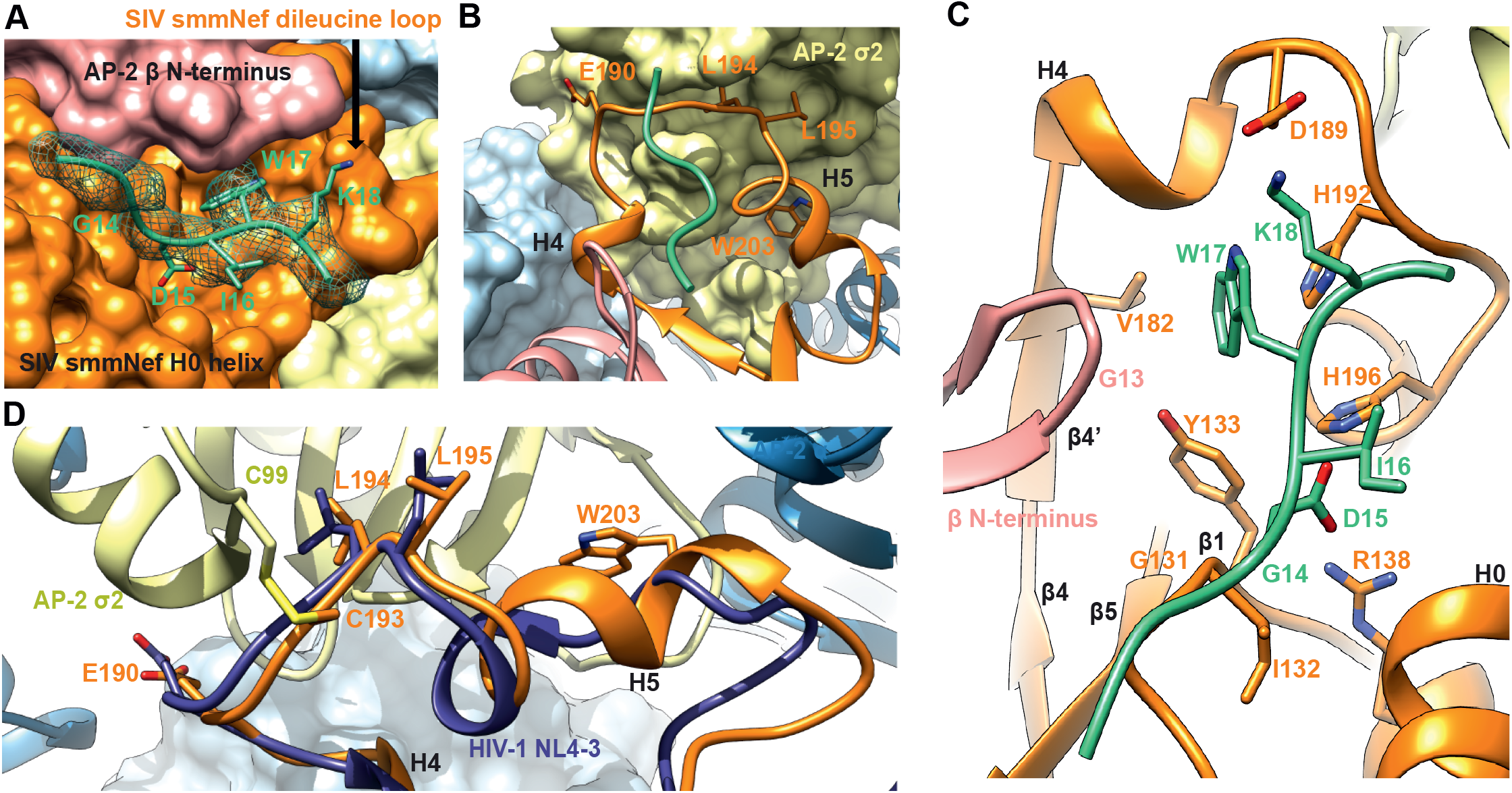
Structural model for cooperative binding of smm tetherin and to SIV smm Nef to AP2^ΔµC^. **(A)** Tetherin GDIWK motif rests in a pocket created by the SIVsmm Nef dileucine loop. The GDIWK motif is further sandwiched by the AP2^ΔµC^ β2 N-terminus and SIVsmm Nef H0 helix. The cryo-EM map density is depicted as a mesh around the smm tetherin peptide. **(B)** Ribbon representation of the SIVsmm Nef dileucine loop bound to the dileucine binding site of the AP2^ΔµC^ α-σ2. Smm tetherin peptide is shown to highlight its position relative to the SIVsmm Nef dileucine loop. **(C)** Hydrophobic pocket or SIVsmm Nef created by both the SIVsmm Nef dileucine loop and Nef core with the smm tetherin GDIWK peptide bound and sandwiched by the AP2^ΔµC^ β2 N-terminus and SIVsmm Nef H0 helix. Interacting residues are highlighted. **(D)** Dileucine loop alignment between SIVsmm Nef of the AP2^ΔµC^:smm tetherin: SIVsmm Nef complex and HIV NL4-3 Nef bound to the AP-2 α-σ2 hemi complex (PDB: 4NEE). Alignment was performed between AP2^ΔµC^ α-σ2 and AP-2 α-σ2 hemi.

### SIVsmm Nef refolds the N-terminus of β2-adaptin

A novel interface was observed between the β2 subunit and smm tetherin (Fig 2A), not found in previous structures of NL4-3 Nef complexes with APs (*21, 25, 27, 28*). Despite a large conformational change upon unlocking, the N-terminus of β2 (residues 13-24) comprises helix α1, preceded by random coil. When AP2^ΔµC^ binds to smm tetherin and SIVsmm Nef, the flexible loop (residues 6-12) along with the first turn of α1 (residues 14-17) refold into a β hairpin structure not previously described in AP structures (Fig. 3A). This new conformation appears to be stabilized by direct interactions with both SIVsmm Nef and the smm tetherin tail (Fig. 3A, B).

**Fig. 3.**
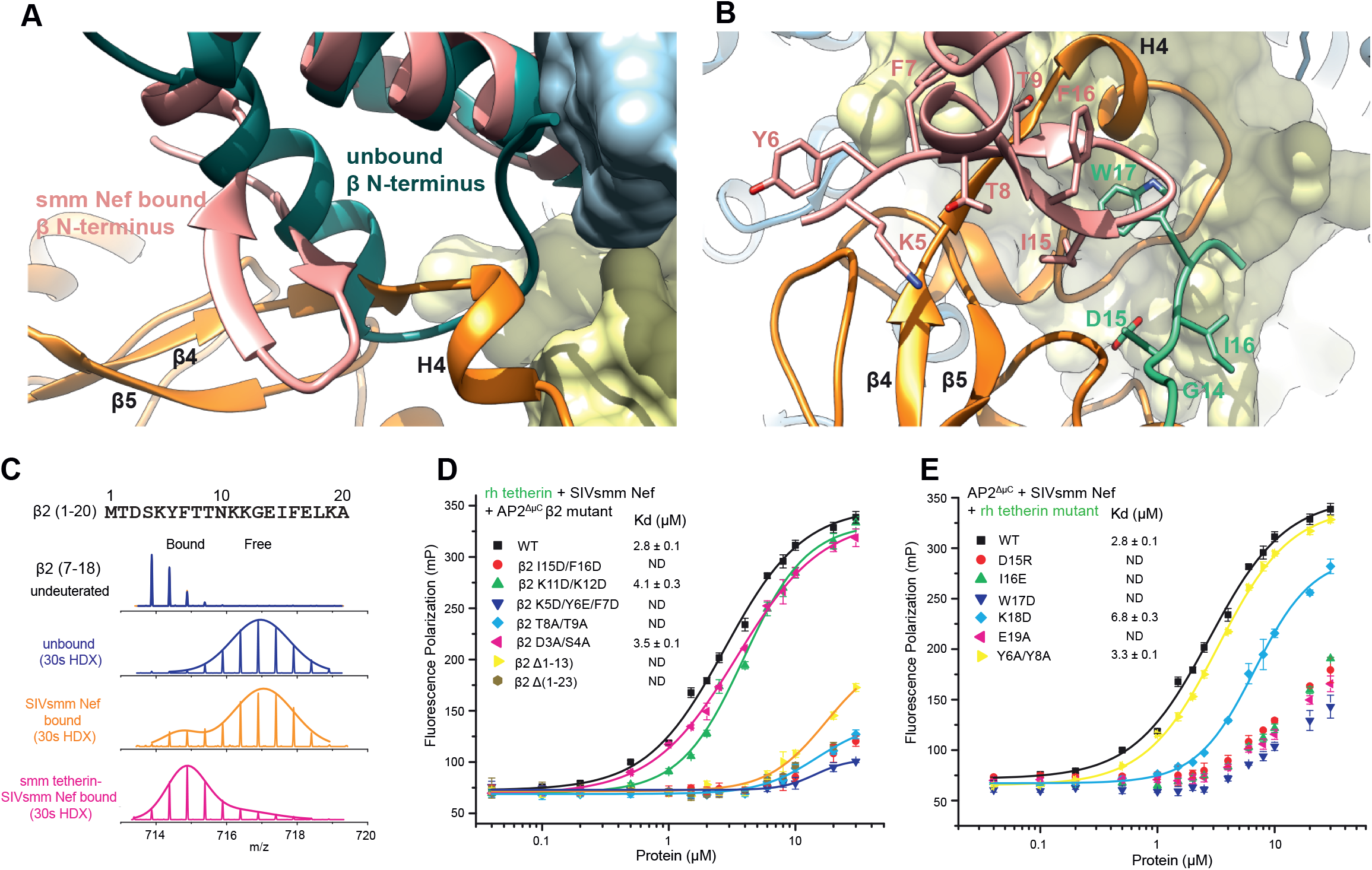
Structural comparison and analysis of the β2-adaptin refolding at the SIVsmm Nef interface. **(A)** Alignment of unlocked AP-2 β2 from PDB:2XA7 onto the AP2^ΔµC^ β2 of the AP2^ΔµC^:smm tetherin: SIVsmm Nef structure. **(B)** Mass spectra of the peptide from β2 (7-18) in AP2^ΔµC^ complex upon SIVsmm Nef (orange) or smm tetherin-SIVsmm Nef (magenta) binding, compared to unbound state (blue). Gaussian fit on one or two peaks is used to represent the distribution of peak heights of the ion peaks across the m/z values and overlay with mass spectra (straight line). **(C)** β2 mutants in β/Nef interface disrupt cooperative tetherin binding, measured by Fluorescence polarization assay. **(D)** FP assay screens the key residues in tetherin tail.

To corroborate the refolding of the β2 N-terminus, we used hydrogen-deuterium exchange coupled to mass spectrometry (HDX-MS) to monitor changes in the dynamics of AP-2 protein upon binding tetherin and SIVsmm Nef. When bound to SIVsmm Nef only, the β2 N-terminus underwent rapid exchange, similar to the unbound state (Fig. S3). However, when bound to both tetherin and Nef, β2 (peptide 7-18) underwent 40% less HD exchange (30 s) (Fig. 3C, S3). Slower deuteration was also seen in α and σ2 residues in SIVsmm Nef binding regions (Fig. S4), as expected. The finding that protection from amide HD exchange in the β2 N-terminus depends on the presence of both SIVsmm Nef and tetherin strongly supports the conclusions drawn from the cryo-EM structure that the presence of these two molecules together is required for refolding of this region.

Side-chains of the induced β2 hairpin present a wall of exposed hydrophobic surface for interactions with tetherin and Nef. These include Tyr6, Phe7, Thr8, Thr9, and Ile15 (Fig. 3B). We generated double and triple mutations to test the role of β2 N-terminal residues in the novel interface (Fig. 3D). Mutation of residues in both the first (residue 5-7 and 8-9) and second β-strands (residue 15-16) disrupted complex formation. Mutation of Lys11 and Lys12 at the turn connecting the two strands had no effect, however, consistent with their high solvent exposure.

SIVsmm Nef residues 179-189 are the ones most intimately involved in interacting with the novel β2 hairpin structure (Fig. 3A, B, Fig. 4A, B). This key section is well-conserved in HIV-1 Nef (Fig. 4A). The SIVsmm Nef main chain from residues 179-182 folds into a β strand that complements the β2 hairpin, completing a three-stranded β-sheet spanning SIVsmm Nef and β2. SIVsmm Nef side-chains of Val180, Ser183, Glu185, and Ala186 all interact with β2 (Fig. 4B). The mutations made to disrupt the function of this region, V180D/S183Y and E185A/A186H, severely reduced binding as judged by FP (Fig. 4C). Others immediately following these, including Gln187, Glu188, Asp189, and Glu190 (the E of the EXXXL(L/V) motif) contact smm tetherin and σ2, and lead into the heart of the dileucine highlighting how this short stretch of Nef sequence forms a central nexus between all of the other elements of the complex (Fig. 4B). The relatively solvent exposed residues 187-188 seem to be non-essential, while mutation of Asp189 or Glu190 have strong reductions in binding (Fig. 4C). Collectively, the cryo-EM, HDX-MS, binding, and mutational data lead to a clear and consistent picture of how tetherin and Nef-induced refolding of the β2 N-terminus creates a novel binding site for the (G/D)DIWK motif in the cytoplasmic tail of simian tetherin.

**Fig. 4.**
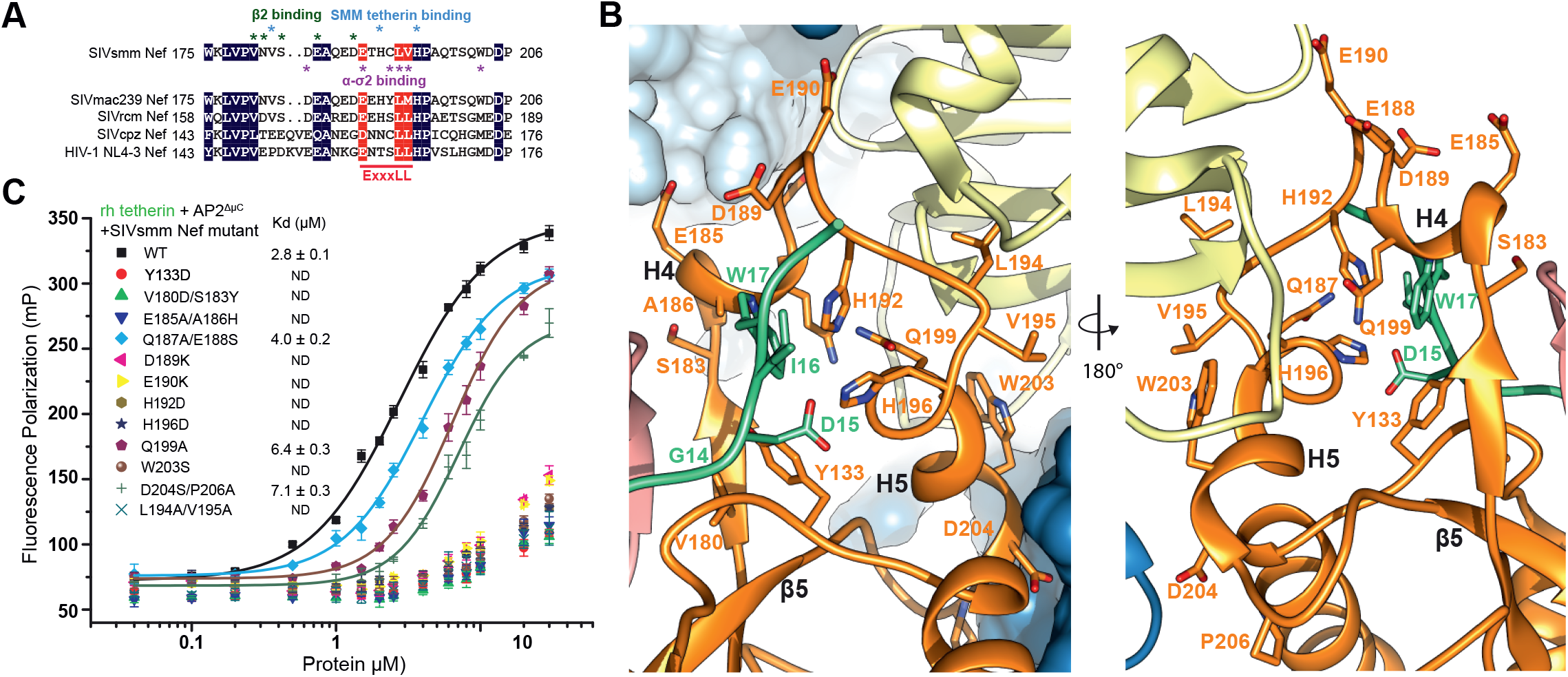
Structural analysis of the SIVsmm Nef tetherin binding pocket. **(A)** Sequence alignment of Nef dileucine loops between SIVsmm Nef and other Nef species. Key residues in SIVsmm Nef for tetherin and AP-2 binding are highlighted. **(B)** Ribbon representation of SIVsmm Nef mutations investigated with smm tetherin GDIW motif shown for reference**. (C)** FP assay indicates that Nef mutations in multi interfaces impair tetherin binding.

### The (G/D)DIWK binding pocket

EM density was resolved for residues 11-19 of smm tetherin (Fig. 2A), with Ile16 and Trp17 serving as landmarks for the register and direction of the tetherin tail. Trp17 is the linchpin of the binding site, and its mutation leads to a severe loss in affinity for the complex (Fig. 3E). Its side-chain is wedged into a pocket between the turn of the AP-2 β2 subunit N-terminal hairpin on the one hand, and SIVsmm Nef strands β1 and β4 and the dileucine loop on the other. The Cα of β2 subunit Gly13 is in direct contact with the tetherin Trp side-chain (Fig. 2C), the closest direct contact between tetherin and AP-2. Tyr133 from Nef strand β1 forms one side of the Trp indole-binding pocket. The remainder of the pocket is formed by residues from Nef strand β4’ and H4, and most critically, the back side of the dileucine loop. Nef strand β4’ contributes Val182 and H4 contributes Ala186. While Nef residues Leu194 and Val195 are the critical plug the connects Nef to the “socket” of the dileucine binding site in σ2, His192 and His196, which point in the opposite direction, are key parts of the Trp binding pocket. The His192 side-chain imidazole stacks against the Trp indole, forming the most extensive interaction of any residue in the pocket.

Of the remaining tetherin side-chains observed, Ile16 forms van der Waals contacts with two large side chains of the Nef N-terminal H0 (Fig. 2C), however the identities of these side-chains could not be assigned with confidence. Density for Asp15 was missing, as often seen in EM density and as seen for most of the Asp and Glu in this reconstruction. On the basis of the surrounding stereochemical constraints, the side-chain was placed such that salt bridges are formed both with His196 of the Nef dileucine loop, and with Arg138 of Nef H2 (Fig. 2C). Mutation of either Asp15 or Ile16 disrupts complex formation (Fig. 3E), consistent with the multiple structural interactions. Single mutations in other residues had little impact on binding, with the exception of Lys18 and Glu19 (Fig. 3E). Lys18 forms a salt bridge with Nef Asp189 (Fig. 2C), which likely accounts for the 2.5-fold reduction in affinity in the K18D mutant. Density for the surface-exposed Glu19 was not visualized, however, so the structural basis for the effect of E19A is unclear. These data are gratifying consistent with biological observations of the critical role of the (G/D)DIWK motif for counteraction by SIV Nef.

### α-σ2 engagement by SIVsmm Nef

The dileucine loop of SIVsmm Nef interacts with the AP-2 α-σ2 hemicomplex in a similar manner to HIV-1 NL4-3 Nef (*21*), but the SIVsmm Nef helix H5 is more regular than its NL4-3 counterpart and has additional interactions. SIVsmm Nef Trp203 anchors the C-terminus of H5 to a pocket formed by residues 60-62 of σ2, which are protected from HDX on complex formation (Fig. S5). The mutation W203S severely impacts complex formation (Fig. 4C). SIVsmm Nef Asp204 and Pro206 also make SIV-specific interactions with σ2 (Fig. 4B). Consistent with the structural observations, HDX-MS showed a very high level of protection for the Nef dileucine loop when the complete ternary complex was formed (Fig. S6). These findings show that SIVsmm Nef is capable for forming a tight and highly ordered complex with α-σ2 *via* the dileucine loop, which is rarely seen in Nef structures not bound to AP-2.

### Residues specifically disrupting anti-tetherin activity of SIVsmm Nef

To examine the relevance of the specific residues for Nef function, we introduced single or combined mutations of Y133D, H192D, H196D, and W203S into the bicistronic pCG expression vector and in proviral HIV-1 NL4-3 constructs coexpressing the SIVsmm Fwr1 Nef protein and eGFP via an IRES element. Western blot analysis showed that most mutant Nef proteins were efficiently expressed (Fig. S7). Exceptions were those containing the Y133D substitution, which showed lower levels of expression than the parental SIVsmm Nef and were thus excluded from functional analyses. To test anti-tetherin activity, we cotransfected HEK293T cells with *env*- and *vpu*-deficient HIV-1 NL4-3 proviral constructs containing wild type or mutant SIVsmm *nef* alleles, as well as smm tetherin and Env expression or control plasmid. Subsequently, we measured tetherin surface levels, infectious virus yield and p24 antigen release two days later. Mutation of H192D or W203S disrupted the ability of SIVsmm Nef to downregulate cell surface expression of smm tetherin (Fig. 5A) and to enhance infectious virus yield (Fig. 5B) and p24 antigen release (Fig. S8). In comparison, H196D did not affect the ability of SIVsmm Nef to downregulate smm tetherin but resulted in a phenotype intermediate between wt and *nef*-defective SIVsmm in infectious virus yield assays.

**Fig. 5.**
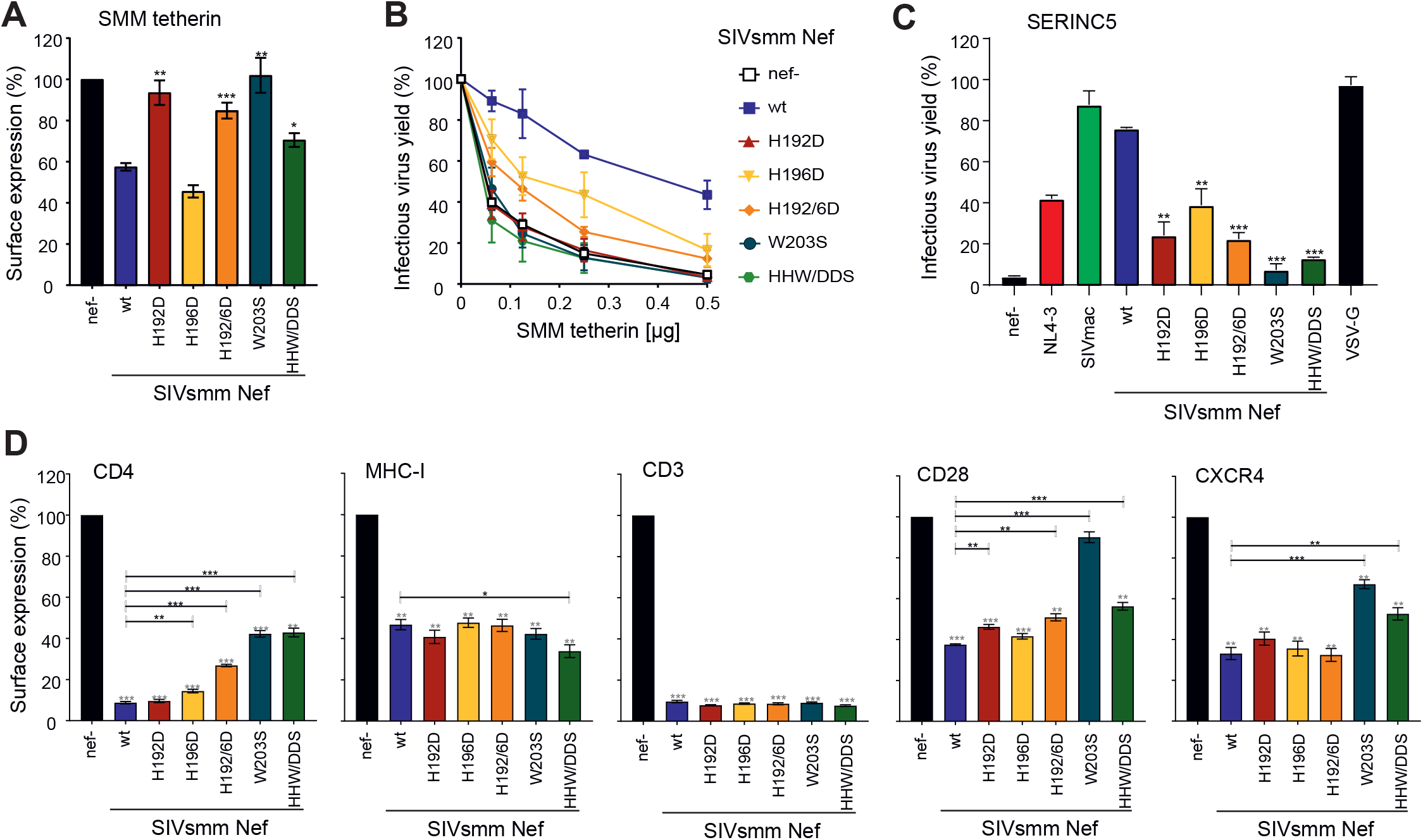
H192D specifically disrupts tetherin antagonism by SIVsmm Nef. **(A, B)** Cell surface expression of tetherin (A) and infectious virus yield (B) from HEK293T cells following co-transfection with an *env*- and *vpu*-deficient HIV-1 NL4-3 proviral construct encoding the indicated *nef* alleles and eGFP via an IRES and varying amounts of plasmids expressing SMM tetherin. (A) To measure tetherin modulation, cells were stained with an APC-conjugated antibody against tetherin (Biolegend, 348410) or an isotype control (mouse IgG1-κ, Biolegend, 400122). Tetherin surface expression was analyzed by flow cytometry and the APC mean fluorescence intensity (MFI) of cells coexpressing Nef and eGFP was normalized to the negative control (*nef*-; 100%). Mean of three to six independent experiments ± SEM. (B) Infectious virus yield was determined by infection of TZM-bl indicator cells and normalized to that detected in the absence of tetherin (100%). Mean of three to six independent experiments measured in triplicates ± SEM. **(C)** HEK293T cells were co-transfected with *vpu-* and *env-*defective HIV-1 NL4-3 constructs encoding the indicated Nef proteins, a construct expressing the HIV-1 Env glycoprotein, and SERINC5 expression or empty control plasmids. Shown are the mean levels of infectious virus production in the presence of transient SERINC5 expression relative to those obtained in cells transfected with the control vector (100%). Mean of three to six independent experiments measured in triplicates ± SEM. **(D)** To measure receptor modulation, human PBMCs were transduced with VSV-G pseudoytped *env*- and *vpu*-deficient HIV-1 NL4-3 IRES-eGFP constructs containing the indicated *nef* alleles. Cell surface expression of infected (i.e. eGFP positive) cells was normalized to uninfected (i.e. eGFP negative) cells within the same sample before normalization to the negative control (*nef*-, 100%). Mean of three independent experiments ± SEM. P-values represent differences from the *nef*-defective (grey stars) or wt SIVsmm (black stars) controls. *, p < 0.05; **, p < 0.01; ***, p < 0.001.

To further analyze the impact of the mutations on SIVsmm Nef function, we examined their effect on viral infectivity in the presence of SERINC5, a recently described restriction factor that impairs virion infectivity (*29, 30*). In contrast to tetherin antagonism, Nef counteracts SERINC5 and modulates various immune receptors in a species-independent manner (*29, 30*). Coexpression of SERINC5 reduced infectivity of the *nef*-defective control HIV-1 construct by >95% but did not affect the infectiousness of VSV-G pseudotyped viral particles (Fig. 5C). The H192D mutation impaired and W203S almost fully disrupted the anti-SERINC5 activity of SIVsmm Nef, while H196D resulted in an intermediate phenotype (Fig. 5C).

To further define the impact of the above-mentioned amino acid changes on SIVsmm Nef function, we transduced human peripheral blood mononuclear cells (PBMCs) with VSV-G pseudotyped *env*- and *vpu*-deficient HIV-1 NL4-3 based IRES-eGFP constructs containing different *nef* alleles and performed flow cytometric analysis to determine surface expression of a variety of cellular receptors (Fig. S9). In agreement with published data (*31*), the parental SIVsmm Nef efficiently downregulated CD4, CD3, and (to a lesser extent) MHC-I, CD28, and CXCR4 in primary human cells (Fig. 5D). Although mutation of H192D fully disrupted the anti-tetherin and anti-SERINC5 activities of SIVsmm Nef it did not reduce its ability to downregulate any of these immune receptors from the cell surface (Figs. 5D, S9). Similarly, mutation of H196D had little if any effect on the efficiency of Nef-mediated modulation of CD3, MHC-I, CD28, and CXCR4 although it moderately attenuated down-modulation of CD4. In comparison, W203S strongly reduced downregulation of CD4 and CXCR4 and fully disrupted the effect on CD28, while having no effect on modulation of MHC-I or CD3 (Fig. 5D). Similar observations on the role of SIVmac Nef residues His196 and Trp203 were recently reported in the context of MHC escape mutants (*32*). Thus, H192 in the Ex<u>x</u>xLL/M motif plays a key role in the ability of Nef to counteract tetherin and SERINC5 but is dispensable for its ability to remove various immune receptors from the cell surface.

## Discussion

It was discovered a decade ago that the sensitivity of simian tetherin to antagonism by SIV Nefs maps to a (G/D)DIWK motif in the N-terminal tail of tetherin that is missing in its human ortholog (*3, 5, 6*). The structure explains how each of the four residues in the most conserved central DIWK part of this motif interact with AP-2 and SIVsmm Nef, and is beautifully congruent with the biological data. The structural and binding data explain how the absence of this motif in human tetherin renders it resistant to downregulation from the cell surface by SIV Nefs. The inability of SIV to antagonize human tetherin is now considered to be one of the major hurdles to successful simian-human zoonotic transmission (*7*). Our data thus provide the structural basis for a transmission barrier that helps to protect humans from efficient primate lentiviral transmission.

The involvement of the AP-2 β2 subunit in recognizing both Nef and cargo sets the SIVsmm Nef and tetherin interaction apart from all other physiological or Nef-hijacked sorting events reported to date. All known physiological cargoes that bind to AP cores do so through either the Tyr or dileucine motif binding sites in the μ and the σ subunits, respectively (*9*). Nef downregulation of MHC-I (*33*) occurs principally through the Tyr binding site on the CTD of μ1. The difference is consistent with the null effect of all of the SIVsmm Nef mutants tested on MHC-I downregulation. CD4 downregulation by Nef-appears to be mediated by a site on Nef wedged between strand β1 and helix H2 (*34*), which is not in direct contact with AP-2. SIV tetherin downregulation shares with CD4, CD28, and CXCR4 a dependence on the engagement of the dileucine loop (*10–12, 35*) with the α-σ2 hemicomplex, consistent with the W203S phenotype in tetherin and CD4 downregulation. It is remarkable that such a small protein as Nef can bind and hijack the AP complexes in so many different ways. This multifunctionality confers robustness to viral mutation and host variation.

Our results suggest that many downregulatory functions of Nef are structurally and genetically separable. Perhaps most notably, mutation of H192D impairs the ability of SIVsmm Nef to antagonize tetherin and SERINC5 during the late stage of the viral replication cycle but had no disruptive effect on the modulation of various immune receptors (CD4, CD3, CD28, MHC-I and CXCR4) required for viral immune evasion throughout the viral life cycle. Thus, this mutation may help probe the relevance of early *vs.* late Nef functions for viral replication and pathogenicity *in vivo* in non-human primate models for AIDS.

His192 is conserved in most monkey SIVs but not the ape and human viruses. On the other hand His196 is conserved across all Nefs, and H196D is defective in SERINC5 downregulation in human cells. The proximity of His196 to the β2 hairpin interface, and the conservation in HIV-1 of the Nef strand β4 involved in hairpin refolding suggests that the β2 hairpin participates in HIV-1 Nef downregulation of SERINC5. The apparent lack of a normal physiological role for the β2 hairpin in AP-2-dependent cargo sorting suggests that this interface could be targeted therapeutically by Nef antagonists to restore SERINC5 expression and block viral infectivity.

Different HIV strains have adapted to their human hosts with different degrees of success through both Nef-dependent and independent mechanisms. HIV-1 O-Nefs adapted to target a tetherin sequence adjacent to the DIWK deletion (*20*) in conjunction with the μ1 subunit of AP-1. When bound to human tetherin and μ1, HIV-1 O-Nef bridges the β1-μ1 and γ1-σ1 hemicomplexes of two different AP-1 tetramers in *trans*. This *trans* bridging mode, in contrast to the *cis* bridge described here for SIVsmm Nef, leads to AP-1 trimerization and formation of a structure capable of sequestering tetherin at the TGN (*21*). HIV-2 Nefs never recovered the ability to target tetherin, and HIV-2 is dependent on Env for tetherin counteraction. The pandemic M-group HIV evolved a potent Vpu counteraction of a tetherin, which however, uses some of the same principles as Nef-dependent counteraction. HIV-1 M-Vpu has a dileucine that can engage the γ1-σ1 hemicomplex of AP-1 and targets the same DIWK deletion-proximal sequence in tetherin and the same site on μ1 as HIV-1 O-Nefs (*13*). All of the adaptions whereby HIVs counteract human tetherin involved major changes in mechanism, either using Nef to hijack a different AP complex through different sites, or using other proteins entirely. This illustrates that the barrier caused by the absence of DIWK in human tetherin cannot be overcome simply by modifying and repurposing the pre-existing DIWK binding site in SIV Nefs. On the other hand, the fact that the barrier ultimately was overcome repeatedly and in different ways highlights the multiplicity of entry points to the host trafficking machinery and the versatility of viral proteins through evolution.

### Ethical statement

Experiments involving human blood were reviewed and approved by the Institutional Review Board (i.e. the Ethics Committee of Ulm University). Individuals and/or their legal guardians provided written informed consent prior to donating blood and all human-derived samples were anonymized before use.

## Methods

### Plasmid construction

The four subunits of AP2^ΔµC^ were subcloned into pST39 vector by restriction cloning (*36*). DNAs coding for the four subunits of AP2^ΔµC^ were as follows: rat σ2 full-length, mouse µ2 (1-141), rat α (1-621) as a C-terminal GST fusion, and rat β2 (1-591) fused to an N-terminal 6xHis tag. TEV cleavage sites were introduced between the affinity tags and the protein. GST tagged AP2^hemi^ consists of rat α (1-396) with an N-terminal GST tag and rat σ2 full-length, the minimal construct binding to HIV-1 NL4-3 Nef (*25*). For all structural and in vitro experiments, the SIVsmm Fwr1 Nef C55A construct bearing the single Cys193 was used, which is referred to as wild-type of SIVsmm Nef. All constructs used in this study are listed in Table S1.

For MBP pull down or binding assay, SIVsmm Nef wild-type and mutants were subcloned into LIC 2BT vector (Macrolab, UC Berkeley), and were expressed as TEV-cleavable N-terminal 6xHis fusions. DNAs coding for the tetherin tail (2-26) from rhesus macaque or sooty mangabey were codon-optimized using DNAworks server (*37*) for *E. coli* expression. Rhesus tetherin was first fused to a C-terminal MBP tag and then subcloned into the LIC 2BT vector, which was used to generate proteins for MBP pull down assays. Rhesus tetherin was also subcloned into pGST2, yielding a fusion with an N-terminal GST and a TEV cleavage site for FP assays (*38*). For cryo-EM, a PCR fragment encoding sooty mangabey tetherin (4-26)-C2S-10aa linker-SIVsmm Nef (1-251)-C55A was subcloned into pMBP2, and the protein was expressed with an N-terminal TEV cleavable MBP tag and C-terminal uncleavable 6xHis tag.

### Protein expression and purification

Wild-type and mutant AP2^ΔµC^ complexes were expressed in BL21 (DE3) pLysS strain (Promega, Madison, WI) by induction at 23 °C overnight. The cells were lysed by sonication in 50 mM Tris pH 8.0, 300 mM NaCl, 10% glycerol, 3 mM β-mercaptoethanol (βME), 20 mM imidazole and 2 mM PMSF. The clarified lysate was first purified on a Ni-NTA column (Qiagen, Valencia, CA). The eluate was further purified on glutathione-Sepharose 4B resin (GE healthcare, Piscataway, NJ). After TEV cleavage at 4 °C overnight, the sample was concentrated and then loaded onto a HiLoad 16/60 Superdex 200 column (GE healthcare) in the sample buffer of 20 mM Tris pH 8.0, 150 mM NaCl, 0.1 mM TCEP. The AP2^ΔµC^ fractions were passed through 1 ml of glutathione-Sepharose 4B resin to capture the GST tag, concentrated, and flash-frozen in liquid N^2^.

His-tagged SIVsmm Nef constructs were expressed in BL21 (DE3) star cells (Life technologies, Grand Island, NY), and induced with 0.3 mM IPTG at 25 °C overnight. Purification was carried out using Ni-NTA resin, and samples were eluted with 200 mM imidazole/500 mM NaCl Tris buffer, pH 8. The eluate was subjected to a HiLoad 16/60 Superdex 75 column in high salt buffer of 20 mM Tris pH 8, 500 mM NaCl, 0.1 mM TCEP.

All MBP tagged proteins were expressed in BL21 (DE3) star cells and grown at 30 °C overnight. Cells were induced at 35 °C for 2 hours. The purification of His-tagged tetherin-MBP proteins was carried out by Ni-NTA column, size-exclusion in a HiLoad 16/60 Superdex 200 column, and then a second Ni-NTA column. Smm tetherin-10aa linker-SIVsmm Nef proteins were first purified using a Ni-NTA column, the eluate was diluted 5 times in Q buffer A (30 mM Tris pH 8), and then loaded onto a HiTrap Q HP 5ml column (GE healthcare, Piscataway, NJ). The elution on Q column was performed with a 10 CV linear gradient from 0-1 M NaCl in Q buffer A. The sample fractions were pooled together for TEV cleavage at 4 °C overnight. The TEV cleaved samples was adjusted to final 20 mM imidazole and 500 mM NaCl, and then passed through the second Ni column to remove free MBP tag. The proteins were eluted in 500 mM NaCl /200 mM imidazole buffer and subjected to a 16/60 Superdex 75 column in the same high salt buffer.

GST-tagged tetherin constructs were expressed in BL21 (DE3) star cells, and induced with 0.3mM IPTG at 20 °C overnight. The cleared lysate was purified using glutathione-Sepharose 4B resin. The eluate was concentrated to 1-1.5 ml, and 150 µl of TEV was added to cleave samples at room temperature for 4-5 hours. The cut proteins were then loaded onto a 16/60 Superdex 75 column in the same sample buffer to separate tetherin peptides from MBP and TEV proteins.

### MBP pull down assay

35 µg of recombinant His-tetherin-MBP proteins were incubated with His-and GST-tagged AP2^ΔµC^ (1 µM) or/and SIVsmm Nef proteins (6 µM) at 4 °C overnight in 20 mM Tris pH 8, 150 mM NaCl, 0.1 mM TCEP. Then 30 µl Amylose resin (New England Biolabs, Ipswich, MA) was to the mixture and rocked at 4 °C for 2 hours. The beads were washed 4 times, mixed with 50 µl of 2x lithium dodecylsulfate (LDS)/ BME buffer and heated at 90°C for 3 min. 19 µl of each sample was subjected to SDS/PAGE gel.

### Tetherin peptide fluorescent labeling

The wild-type of rh tetherin peptide used in the FP assay consists of residue (2-26): APILYDYRKMPMGDIWKEDGDKRCK. The single cysteine at the C-terminus was used for labeling. The rh tetherin tail differs by only two residues from the smm tetherin tail, which are both outside the binding region. Alexa Fluor 488 C^5^ Maleimide (Thermo Fisher Scientific, Waltham, MA) was dissolved in dimethyl sulfoxide (DMSO). 100 µM of diluted peptide in 1x PBS pH 7.4 was reacted with dye at 1:1.2 molar ratio for 40 min at room temperature. The conjugation reaction was quenched with 100 mM βME at 15 min at room temperature. The samples were then subjected to a HiTrap desalting column in the sample buffer. The Alexa488 labeling efficiency of each peptide was between 96% and 100%.

### Fluorescence polarization (FP) binding assay

Assays were conducted in Greiner 384-well black microplates (784076, Greiner, Monroe, NC). SIVsmm Nef proteins were exchanged from high salt buffer to the sample buffer using a Zeba spin desalting column (7K MWCO, Thermo Fisher Scientific). After desalting, the stock concentration of SIVsmm Nef was adjusted to 80 µM. The Alexa488 labeled tetherin tail peptide was incubated with SIVsmm Nef and AP2^ΔµC^ in a total volume of 20 µl for 30 min at room temperature. The final concentration of labeled peptide was fixed at 50 nM, while SIVsmm Nef and AP2^ΔµC^ were kept at a 1:1 molar ratio and titrated from 0.04 to 30 µM in this assay. FP values (mP) were measured using a Synergy H4 microplate reader (excitation at 485 nm and emission at 528 nm, BioTek, Winooski, VT), and plotted as a function of protein concentration using a logarithmic scale (*39, 40*). Measurements were repeated at three times and the data were processed using Origin (OriginLab, Northampton, MA). The binding constant (*K*d) was fitted using a one-site model.

### smm tetherin-SIVsmm Nef: AP2^ΔµC^ cross-linking

In order to prevent dissociation of the AP-2:Nef complex, intermolecular disulfide cross linking activated by 2,2’-Dipyridyldisulfide (2-PDS) was used to stabilize the SIVsmm Nef-AP-2 interaction. Ser163 in the Nef dileucine loop is within 3Å of Asn97 in σ2 (PDB: 4NEE) (*25*). The equivalent residue in SIVsmm Nef is Cys193, thus we generated the σ2 N97C/C99S mutation (denoted AP2^ΔµC^SS following cross-linking) to induce this intermolecular disulfide bond formation. AP2^ΔµC^SS proteins were treated with final 20 mM DTT at 30 °C for 30min, followed by desalting into cross-linking buffer (20 mM Tris pH 8, 150 mM NaCl) using Zeba spin desalting column. smm tetherin-SIVsmm Nef fusion proteins were also reduced by 20 mM DTT at 30 °C for 30min, followed by desalting into the buffer of 20 mM Tris pH 8, 300 mM NaCl using Zeba spin column. After elution from the desalting column, tetherin-Nef fusion proteins were treated with final 3 mM 2,2’-Dipyridyldisulfide (2-PDS), and incubated at room temperature for 30min (*41*). The activated tetherin-Nef samples were desalted into 20mM Tris PH 8, 150mM NaCl to remove excess 2,2’-Dipyridyldisulfide. Then the activated samples (final

1.5 µM) were immediately mixed with desalted AP2^ΔµC^ss (final 1 µM) and incubated at RT for 30min. The cross-linking mixture was further purified over a HiTrap Q HP 1ml column, and then subjected to a Superdex 200 10/300 GL column in cross-linking buffer. The AP2^ΔµC^SS:smm tetherin (4-26)-C2S-10aa-SIVsmm Nef (1-251)-C55A complex eluted as a single homogenous peak in gel filtration, and no additional products were observed on SDS-PAGE.

### Cryo-EM Data Collection

AP2^ΔµC^SS:smm tetherin (4-26)-C2S-10aa-SIVsmm Nef (1-251)-C55A samples for data collection were prepared on 2/1-thick C-flat holey carbon copper 300-mesh grids that were prepared by plasma cleaning for 30 s at 25 mA using a Pelco easiGlow glow discharger (Ted Pella, Inc.). Complex at 0.6 mgmL-1 was applied in a 4 μL drop to one side of the grid, followed by a 30 second wait time, then plunge frozen into liquid ethane using a Vitrobot mark VI. The humidity was controlled at 100%, 22°C and the grids blotted with Whatman #1 paper using a blot force of 6 for 5 s. Two rounds of data collected (dataset 1 and 2) were performed on a Titan Krios (FEI; BACEM UCB) at a nominal magnification of 105,000 and a magnified pixel size of 1.149 Å pix-1. The dose rate was 6.08 e-/Å^2^/sec at the sample. Data collection was carried out using SerialEM. Dataset 1, consisting of 1647 movies, used stage shift navigation with 3 exposures per hole and focusing for each hole. Dataset 2 consisted of 10,248 micrographs using image shift navigation. 3 shots per hole were collected in a 3×3 grid of holes, focusing once per 9 holes. The defocus range was 1.00 to 2.5μm. Movies were acquired on a K2 direct electron detector (Gatan) in super-resolution counting mode with a dose fractionated frame rate of 250 ms and total collection time of 8250 ms.

### Cryo-EM Data Processing

Figure S2 shows the cryo-EM data processing workflow. In brief, gain-corrected movies from two data sets (dataset 1 and 2) were motion corrected using the Relion-3 (*42*) MotionCor2 (*43*) wrapper. Corrected micrographs were inspected visually and micrographs with apparent defects excluded from further processing. The contrast transfer function (CTF) of non-dose weighted micrographs was estimated using Gctf (*44*) for dataset 1 and CTFfind4 (*45*) for the dataset 2. After an additional visual cleanup, 622,487 and 5,696,434 particles were picked from 1618 and 9333 micrographs using Gautomatch and Relion Laplacian particle picking for datasets 1 and 2, respectively. Particles were extracted from datasets 1 and 2 with a 256 pixel box size from dose-weighted micrographs and subjected to a single round of 2D classification at 4-fold binning in Relion-3.0 (*46, 47*) and cryoSPARC v2 (*48*), respectively. Obvious “Junk” was removed resulting in 4,107,380 refined particles. A 4 class Ab-initio 3D reconstruction was performed in cryoSPARC v2 on dataset 1, resulting in 2 heterogeneous 3D classes (class 1 and 2), where class 1 was the object of refinement, and two “junk” classes. Using 3D class 1 and class 2, multiple rounds of heterogeneous 3D refinement were performed on dataset 1 resulting in a set of 215,685 particles in class 1 which refined to 4.25 Å resolution. These 215,685 particles were then transferred to Relion-3.0. When needed, cryoSPARC v2 database files were converted using UCSF PyEM (https://github.com/asarnow/pyem) and in-house written scripts (https://github.com/simonfromm/miscEM) into star files for the use as inputs in Relion3.0. A single round of 3D classification was run against two low resolution maps simulated from a refined coordinate model built into the original 4.25 Å structure, one containing Nef and the N-terminus of the β2 subunit of AP2^ΔµC^SS (class 1) and one lacking both (class 2). This focused 3D classification was run with a low pass filter of 20 Å, τ = 20, and no angular refinement. 90,285 particles were classified to generate the map containing Nef and the N-terminus of the β2 subunit of AP2^ΔµC^. After further refinement in cryoSPARC v2, final map contained 84,834 particles and refined to 3.7 Å.

The refined particles of dataset 2 were processed using an iterative 2D classification and heterogeneous refinement workflow similarly to dataset 1 (Fig S2). Eventually, 300,243 particles were subject to a round of focused 3D classification in Relion-3.0 identical to the one performed on dataset 1. The resulting 129,701 particles in class 1 were retained and used for a 3D auto-refine run in Relion-3.0 which served as the basis for per-particle CTF refinement and Bayesian polishing in Relion-3.0. After further refinement of the “shiny” particles in cryoSPARC v2, the remaining 128,842 particles of were combined with the 84,834 particles remaining from dataset 1 to give a final dataset of 212,541 particles. Two further rounds of 2D classification and selection yielded 159,406 particles. A final round of homogeneous as well as non-uniform refinement was performed separately in cryoSPARC v2 (Fig S2, S3B). The Non-uniform reconstructions reached a resolution of 3.8 Å and the homogeneous reconstruction reached a resolution of 3.9 Å, both according to the FSC 0.143 cutoff. Both maps showed improved density in the regions of primary interest compared to the prior 3.7 Å map reconstruction. After visual inspection of both reconstructions, the map resulting from non-uniform refinement was chosen for atomic model building and coordinate refinement, and density thereof is shown throughout the manuscript.

### Model Building and Refinement

Rigid body fitting was performed in UCSF Chimera (*49*) using as starting models PDB entries 4NEE and 2XA7, followed by manual adjustment in Coot (*50*). To help alleviate local differences in map quality, two differently filtered and sharpened maps were used during the model building process. Model building in Coot and real-space refinement in phenix (*51*) was used to improve the model geometry against both a locally filtered and b-factor sharpened map generated through the cryoSPARC v2 filtering/sharpening routine (applied b-factor: −145.0 Å^2^) and a map generated from the Relion3.0 PostProcess routine (applied b-factor: −77.4 Å^2^; low-pass filter: 3.8 Å). Real-space refinement as implemented in phenix with a target resolution of 3.8 Å, secondary structure restraints and a weight of 1 was used iteratively with manual adjustments in coot to refine the atomic coordinate model. EMRinger scores were calculated (*52*) to identify regions requiring further remodeling in Coot. Where clear residue side-chain density was lacking, side chains were not modeled.

The final model-map cross correlation was 0.75, with a map-versus-model FSC0.5 of 4.5 Å (Fig S3D). Model geometry was validated using MolProbity (*53*). All map and model statistics are detailed in Table S2. A cross-validation test was performed to assess overfitting. In brief, atoms of the final coordinate model were perturbed by an average of 0.5 Å and one round of automated real-space-refinement in phenix against one half map of the gold-standard refinement was performed. The resulting map-vs-model FSC curves of the refined model against the same half map (FSCwork) and the second half map (FSCtest) do not diverge significantly indicating the absence of overfitting (Fig. S3E). Table S2 and Figure S3F summarize the completely assembled AP2^ΔµC^SS:smm tetherin-SIVsmm Nef coordinate model, with S3F showing representative density for all AP-2 subunits, SIV smmNef and smm tetherin.

### HDX-MS

Uncrosslinked SIVsmm Nef-ΔL1-6His was used in all HDX studies. The construct used for HDX contained a deletion of residues 28-36, which are non-conserved in SIVcpz Nef or HIV-1 NL4-3 Nef, and whose presence reduced the stability of the complex. AP2^ΔµC^ was incubated with smm tetherin-SIVsmm Nef-ΔL1 or 10aa-SIVsmm Nef-ΔL1 at 4 °C overnight. The mixture was purified using Ni-NTA resin, and loaded on a Suerpdex 200 10/300 GL column. Fractions were pooled and concentrated to 20 µM. Amide HDX-MS was initiated by a 20-fold dilution of 20 µM sample stock into a D^2^O buffer containing 20 mM HEPES (pD 7.2), 200 mM NaCl and 0.5 mM TCEP at room temperature. After intervals of 30 s-120 s, exchange was quenched at 0 °C with the addition of ice-cold quench buffer (400 mM KH^2^PO^4^/H^3^PO^4^, pH 2.2). The samples were then injected onto an HPLC (Agilent 1100) with in-line peptic digestion and desalting. Desalted peptides were eluted and directly analyzed by an Orbitrap Discovery mass spectrometer (Thermo Scientific). Peptides were identified with tandem MS/MS experiments and a Proteome Discoverer 2.1 (Thermo Scientific) search. Mass analysis of peptide centroids was performed using HDExaminer (Sierra Analytics, Modesto, CA), followed by manual verification of each peptide. The relative deuteron content of the peptic peptides was determined from the centroid of the molecular ion isotope envelope.

### Proviral constructs

Overlap-extension PCR was used to replace the NL4-3 *nef* in the IRES-eGFP HIV-1 construct by SIVsmm *nef* genes as described (*26*). The integrity of all PCR-derived inserts was confirmed by sequencing. The control HIV-1 NL4-3 *env* Δvpu* IRES-eGFP constructs expressing the NL4-3 and SIVmac239 Nefs or containing a disrupted *nef* gene have been previously described (*54*).

### Expression vectors

*Nef* genes were amplified by PCR with flanking primers introducing XbaI and MluI restriction sites for cloning into the bi-cistronic CMV promoter-based pCG IRES eGFP vector as described (*54*). pCG SMM tetherin and pBJ6 human SERINC5-HA protein expression vectors have been previously described (*55, 56*).

### Cell lines

Human Embryonic Kidney (HEK) 293T cells (*57*) were obtained from the American Type Culture Collection (ATCC) and TZM-bl reporter cells (*58*) were kindly provided by Drs. Kappes and Wu and Tranzyme Inc. through the NIH AIDS Reagent Program. Both cell lines were cultured in Dulbecco’s Modified Eagle Medium (DMEM) supplemented with 10% heat-inactivated fetal calf serum (FCS), 2 mM L-glutamine, 100 units/ml penicillin and 100 μg/ml streptomycin. TZM-bl cells express CD4, CCR5 and CXCR4 and contain the β-galactosidase gene under the control of the HIV-1 promoter.

### Primary human cells

Peripheral blood mononuclear cells (PBMCs) from healthy human donors were isolated using lymphocyte separation medium (Biocoll separating solution; Biochrom), stimulated for 3 days with phytohemagglutinin (PHA; 2 μg/ml), and cultured in RPMI 1640 medium with 10% fetal calf serum and 10 ng/ml interleukin-2 (IL-2).

### Western blots

To examine the expression of SIVsmm Nef proteins, HEK293T cells were transfected in 12-well dishes with 2.5 μg DNA of pCG IRES eGFP vectors expressing AU1-tagged Nefs. Two days post-transfection, cells were lysed and proteins separated, blotted and detected as previously described (*56*).

### Flow cytometry

CD4, MHC-I, CD28, CD3, CXCR4 and tetherin surface levels and GFP reporter molecules in PBMCs were measured as described previously (*59*). To analyze cell surface receptor modulation in primary cells, PBMCs were infected with VSV-G-pseudotyped HIV-1 NL4-3 IRES eGFP expressing different *nef* alleles. Three days post infection, cells were stained for CD4 (MHCD0405, Life Technologies), CD3 (555333, BD), MHCI (R7000, Dako), CD28 (348047, BD), or CXCR4 (555976, BD) and analyzed by fluorescence-activated cell sorting (FACS). To calculate the specific effect of Nef on receptor cell surface expression, the mean fluorescence intensities obtained for cells coexpressing Nef and GFP were normalized to those obtained with the *nef*\* construct expressing GFP only (100%).

### Virus stocks

To generate virus stocks, HEK293T cells were co-transfected with the proviral HIV-1 NL4-3 constructs encoding containing various *nef* alleles and a plasmid (pHIT-G) expressing VSV-G to achieve comparably high infection levels for flow cytometric analysis and replication kinetics. Two days post-transfection, supernatants containing infectious virus were harvested. The amount of HIV-1 capsid protein was quantified by p24 antigen ELISA for normalization of the virus dose.

### Viral infectivity

Virus infectivity was determined using TZM-bl cells. Briefly, the cells were sown out in 96-well dishes in a volume of 100 μl and infected with virus stocks produced by transfected HEK293T cells. Three days post-infection, viral infectivity was detected using the Gal-Screen kit from Applied Biosystems as recommended by the manufacturer. β-galactosidase activity was quantified as relative light units per second using the Orion microplate luminometer.

### SERINC5 antagonism

To measure Nef-mediated SERINC5 counteraction, HEK293T cells were co-transfected using calcium phosphate with 1.5 μg of HIV-1 NL4-3 *env* Δvpu* IRES GFP reporter proviral constructs containing various *nef* alleles, 0.05µg pCAGGS_NL4-3 FLAG-tag env or 0.25µg pHIT VSV-g expression plasmid, and 1.25 μg pBJ6-SERINC5-HA expression plasmid or pBJ6-empty vector (12-well format). The HIV-1 NL4-3 *env* Δvpu nef-* construct was used as a negative control, whereas HIV-1 NL4-3 *env* Δvpu nef-* pseudotyped with VSV-G as well as HIV-1 NL43 *env* Δvpu nef+* IRES eGFP and HIV-1 NL43 *env* Δvpu* SIVmac239 *nef* IRES eGFP served as positive controls. Two days post-transfection cell supernatants were harvested and infectious HIV-1 yield was quantified by TZM-bl infection assay. Yields obtained in the presence of pBJ6 SERINC5-HA were normalized to the corresponding pBJ6 empty vector controls (100%) for each virus.

### Tetherin antagonism

To measure Nef-mediated SMM tetherin counteraction, HEK293T cells were co-transfected using calcium phosphate method with 4 μg HIV-1 NL4-3 provirus lacking Vpu expression (Δvpu), 1 μg Nef expression vector or empty vector as well as increasing amounts of SMM tetherin DNA. Two days post-transfection, supernatants and cells were harvested. Infectious HIV-1 yield was quantified by a 96-well TZM-bl infection assay whereas p24 concentration was determined by p24 ELISA quantification of cell free (CF) and cell associated (CA) capsid p24 antigen. Release was calculated as the percentage of CF p24 out of total (CF + CA) produced p24 antigen.

### Statistical methods

The mean activities were compared using Student’s t-test. Similar results were obtained with the Mann Whitney U test. The software package StatView version 4.0 (Abacus Concepts, Berkeley, CA) was used for all calculations.

## Supporting information

Combined SI

## Acknowledgments

We thank D. Toso, S. Fromm, and A. Yokom for cryo-EM support and advice and D. Sauter for comments on the manuscript. Access to the FEI Titan Krios was provided through the BACEM UCB facility. This research was supported by NIH grants R01 AI120691 (X. R.), P50 GM082250 (J. H. H.), F32 GM125209 (C. Z. B.) and an Advanced ERC grant ‘Anti-Virome’ and DFG-funded SFB 1279 and SPP 1923 (F. K.). Additional support was provided by a generous gift from the Andrew Dougherty Vision Foundation (C.Z.B.) EM density has been deposited in the EMDB with accession code EMD-20217. Atomic coordinates have been deposited in the PDB with the accession code 6OWT.

## Author Contributions

J.H.H. and X.R. conceived the overall project. C.Z.B., X.R., C.M.S., E.H., and D.K. performed research. J.H.H., X.R. and F.K. supervised research. J.H.H., C.Z.B., X.R. and F.K. wrote the initial manuscript and all participated in editing.

