## Supplementary material for "Structural basis for tetherin antagonism as a barrier to zoonotic lentiviral transmission": Combined SI

**Supplementary Figure Legends**

**Fig S1. AP2$^{\Delta\mu C}$SS:smm tetherin-SIVsmm Nef complex cryo-EM data, 2D class averages and orientation distribution. (A)** Exemplary cryo-EM micrograph at -2.2 μm defocus. Scale bar 500 Å. **(B)** Power spectrum and CTF estimation of the micrograph shown in (A). White line corresponds to 1/3 Å-1 frequency. **(C)** 2D class averages of the AP2$^{\Delta\mu C}$SS:smm tetherin-SIVsmm Nef data. Scale bar is at 200 Å. (D) Orientation distribution of the aligned AP2$^{\Delta\mu C}$SS:smm tetherin: SIV smm Nef particles.

**Fig S2. AP2$^{\Delta\mu C}$SS:smm tetherin-SIVsmm Nef complex cryo-EM data processing workflow.** Software used in the respective step is indicated by italic font.

**Fig S3. AP2$^{\Delta\mu C}$SS:smm tetherin-SIVsmm Nef resolution estimation, atomic coordinate model building and validation. (A)** Fourier shell correlation (FSC) of the final non-uniform refinement with the 0.143 cutoff indicated. **(B)** Comparison between FSC curves for non-uniform and heterogeneous refinement of the final. **(C)** AP2$^{\Delta\mu C}$SS:smm tetherin-SIVsmm Nef map color-coded by the local resolution estimation. **(D)** Non-uniform refinement (teal, map) and map-vs-model FSC (purple, model). The 0.143 (map) and 0.5 (model) cutoffs are indicated. **(E)** Cross-validation test FSC curves to assess overfitting. The refinement target resolution (3.8 Å) is indicated. **(F)** Exemplary fits of the refined model in the reconstructed cryo-EM density.

**Fig S4. Mapping binding sites of β2 in AP2$^{\Delta\mu C}$ complex by HDX-MS. (A)** The HDX difference map of β2 (1-591) in AP2$^{\Delta\mu C}$ complex between smm tetherin-SIVsmm Nef binding and SIVsmm Nef binding at 30, 60, 120 s. **(B)** The HDX difference map of β2 (1-591) in AP2$^{\Delta\mu C}$ complex between smm tetherin-SIVsmm Nef binding and unbound state at 30, 60, 120 s. The percentage changes are color-coded according to the scale bar.

**Fig S5. Mapping binding sites of α and σ2 in AP2$^{\Delta\mu C}$ complex by HDX-MS. (A)** The HDX difference map of α (1-621) in AP2$^{\Delta\mu C}$ complex between smm tetherin-SIVsmm Nef binding and unbound state at 30, 60, 120 s. **(B)** The HDX difference map of σ2 in AP2$^{\Delta\mu C}$ complex between smm tetherin-SIVsmm Nef binding and unbound state at 30, 60, 120 s. The percentage changes are color-coded according to the scale bar.

**Fig S6. Mapping binding sites of SIVsmm Nef by HDX-MS. (A)** Schematic representation of SIVsmm Nef domain structure. Regions interacting with tetherin or AP-2 in the structure are delineated. **(B)** Difference plot of percentage of deuteron incorporation in each residue of SIVsmm Nef binding to $AP2^{\Delta\mu C}$ vs unbound state, 60 s in D2O. **(C)** Difference plot of percentage of deuteron incorporation in each residue of SIVsmm Nef binding to $AP2^{\Delta\mu C}$: smm tetherin vs deuterons incorporated in presence of $AP2^{\Delta\mu C}$, 60 s in D2O. The Δ%D is color-coded according to the scale bar (Inset).

**Fig S7. Expression of the parental and mutant SIVsmm Nef proteins.** Western blot analysis of cell lysates following transfection of HEK293T cells with plasmids expressing AU1-tagged versions of the indicated Nef proteins as well as eGFP via an IRES. eGFP is shown as loading and transfection control.

**Fig S8. Anti-tetherin activity of SIVsmm Nef proteins.** Release of p24 capsid antigen from HEK293T cells following co-transfection with an *env*- and *vpu*-deficient HIV-1 NL4-3 proviral construct expressing the indicated *nef* alleles and eGFP via an IRES and varying amounts of plasmids expressing SMM tetherin. Cell-free and cell-associated p24 was measured by ELISA. Left panel: p24 release was calculated as the amount of p24 in the supernatant relative to total p24 (p24 release = $p24_{supernatant}/(p24_{supernatant} + p24_{cells})$) and is shown as percentages of that detected in the absence of tetherin (100%). Mean of three to six independent experiments ± SEM. Right panel: The area under the curve (AUC) was calculated for each experiment and normalized as n-fold of the negative control (*nef*-, 1). Mean of three to six independent experiments ± SEM.

**Fig S9. Modulation of cell surface receptors by SIVsmm Nef proteins.** Flow cytometric analysis of CD4, MHC-I, CD3, CD28, and CXCR4 surface expression in PBMCs transduced with VSV-G pseudotyped *env*- and *vpu*-deficient HIV-1 M NL4-3 IRES eGFP recombinants expressing the indicated *nef* alleles and eGFP via an IRES. Shown are examples for primary data with the gating strategy and the MFI of the stained surface receptor. Cell surface expression of infected (i.e. eGFP positive) cells was normalized to uninfected (i.e. eGFP negative) cells within the same sample before normalization to the negative control (*nef*-, 100%).

**Movie S1. AP2$^{\Delta\mu C}$SS:smm tetherin-SIVsmm Nef reconstruction.** AP2$^{\Delta\mu C}$SS:smm tetherin-SIVsmm Nef density fit, dileucine binding site, GDIWK binding in the hydrophobic pocket and overall density of the GDIWK motif are highlighted.

**Movie S. β2-adaptin refolding at the SIVsmm Nef interface.** Smm tetherin bound SIVmm Nef binding to AP-2 α-σ2, β2-adaptin refolding at the SIVsmm Nef interface and formation of a β hairpin are highlighted.

### Figure S1

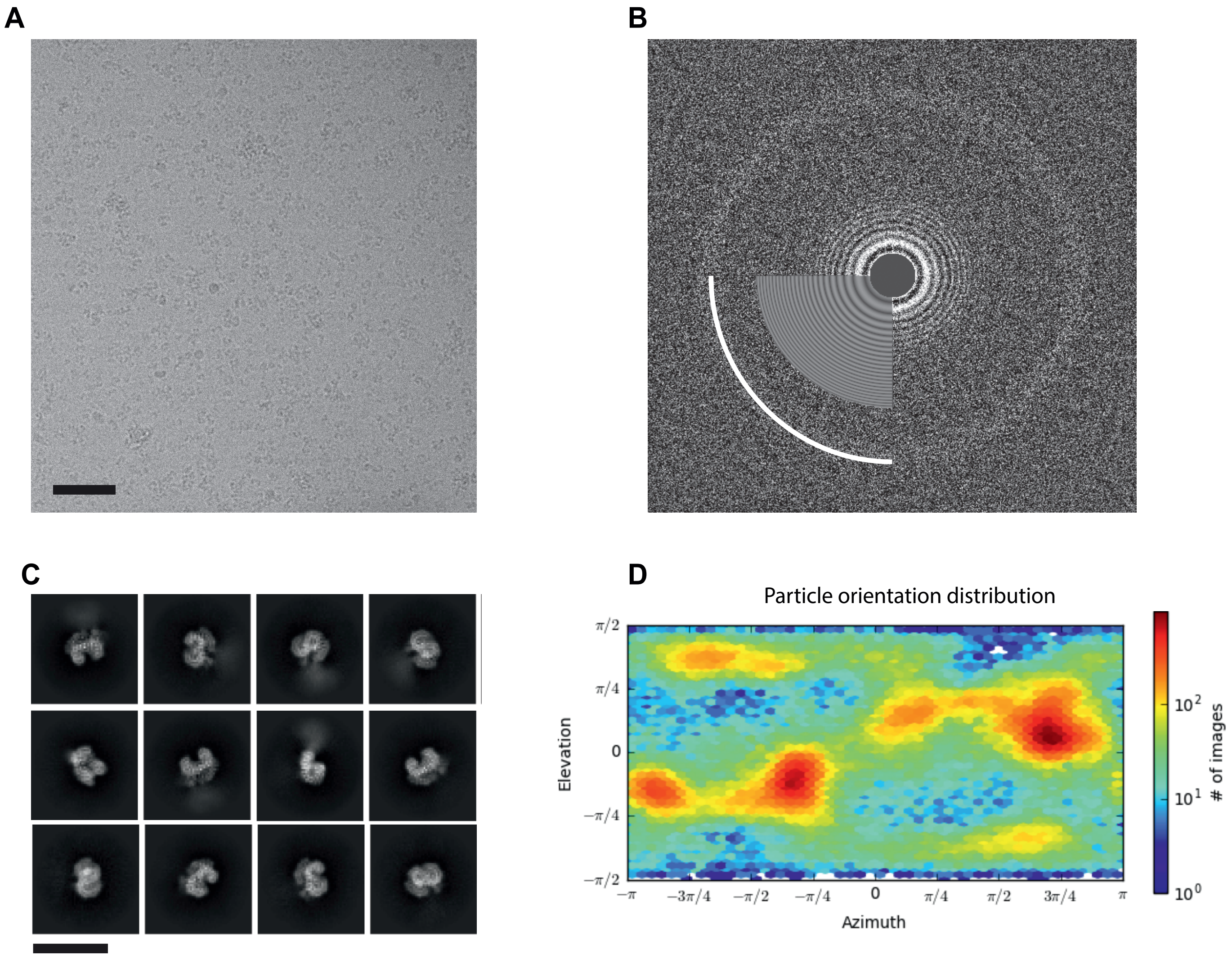
A
B
C
D
Particle orientation distribution
π/2
π/4
0
−π/4
−π/2
Elevation
−π
−3π/4
−π/2
−π/4
0
π/4
π/2
3π/4
π
Azimuth
10^2
10^1
10^0
### of images

### Figure S2

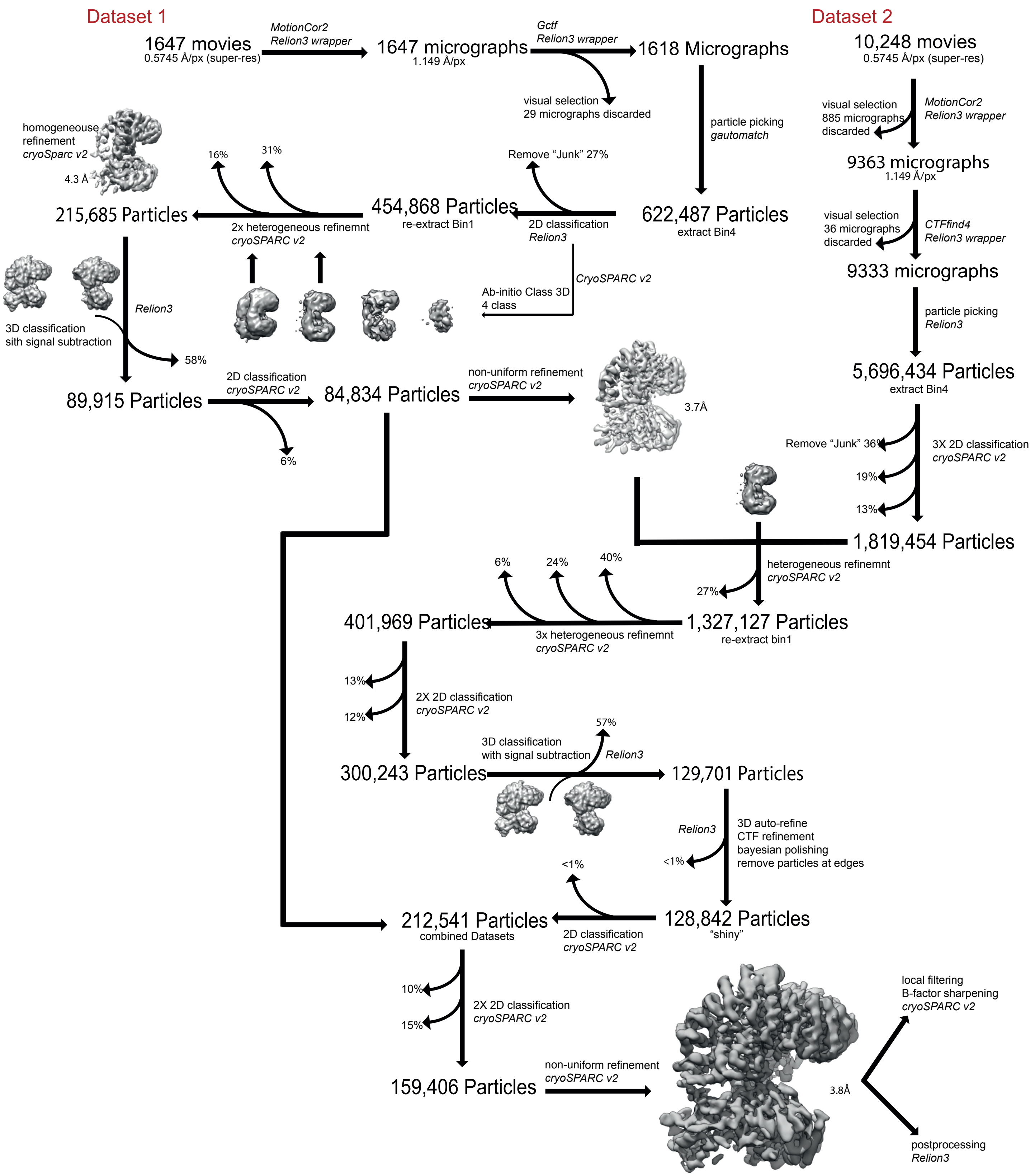

Dataset 1
1647 movies
0.5745 Å/px (super-res)
MotionCor2
Relion3 wrapper
1647 micrographs
1.149 Å/px
Gctf
Relion3 wrapper
1618 Micrographs
visual selection
29 micrographs discarded
particle picking
gautomatch
622,487 Particles
extract Bin4
Remove "Junk" 27%
2D classification
Relion3
454,868 Particles
re-extract Bin1
16%
31%
2x heterogeneous refinemnt
cryoSPARC v2
215,685 Particles
homogeneouse
refinement
cryoSparc v2
4.3 Å
CryoSPARC v2
Ab-initio Class 3D
4 class
Relion3
3D classification
sith signal subtraction
58%
89,915 Particles
2D classification
cryoSPARC v2
6%
84,834 Particles
non-uniform refinement
cryoSPARC v2
3.7Å
Dataset 2
10,248 movies
0.5745 Å/px (super-res)
visual selection
885 micrographs
discarded
MotionCor2
Relion3 wrapper
9363 micrographs
1.149 Å/px
visual selection
36 micrographs
discarded
CTFfind4
Relion3 wrapper
9333 micrographs
particle picking
Relion3
5,696,434 Particles
extract Bin4
Remove "Junk" 36%
3X 2D classification
cryoSPARC v2
19%
13%
1,819,454 Particles
heterogeneous refinemnt
cryoSPARC v2
27%
6%
24%
40%
401,969 Particles
3x heterogeneous refinemnt
cryoSPARC v2
1,327,127 Particles
re-extract bin1
13%
12%
2X 2D classification
cryoSPARC v2
300,243 Particles
3D classification
with signal subtraction
Relion3
57%
129,701 Particles
Relion3
<1%
3D auto-refine
CTF refinement
bayesian polishing
remove particles at edges
<1%
212,541 Particles
combined Datasets
2D classification
cryoSPARC v2
128,842 Particles
"shiny"
10%
15%
2X 2D classification
cryoSPARC v2
159,406 Particles
non-uniform refinement
cryoSPARC v2
3.8Å
local filtering
B-factor sharpening
cryoSPARC v2
postprocessing
Relion3

### Figure S3

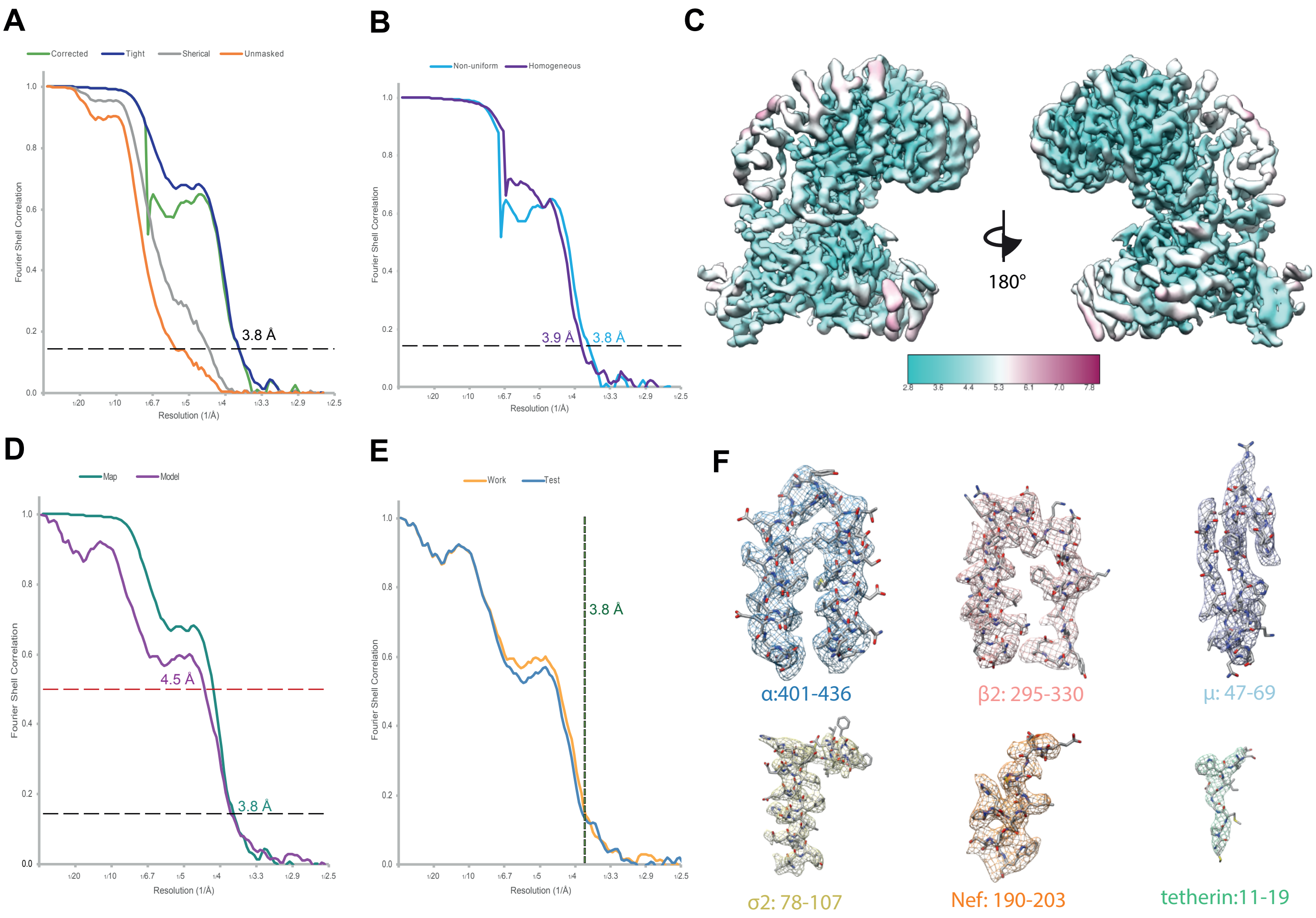

A
Corrected
Tight
Sherical
Unmasked
Fourier Shell Correlation
Resolution (1/Å)
3.8 Å
B
Non-uniform
Homogeneous
3.9 Å
3.8 Å
C
180°
D
Map
Model
4.5 Å
3.8 Å
E
Work
Test
3.8 Å
F
α:401-436
β2: 295-330
μ: 47-69
σ2: 78-107
Nef: 190-203
tetherin:11-19

### Figure S4

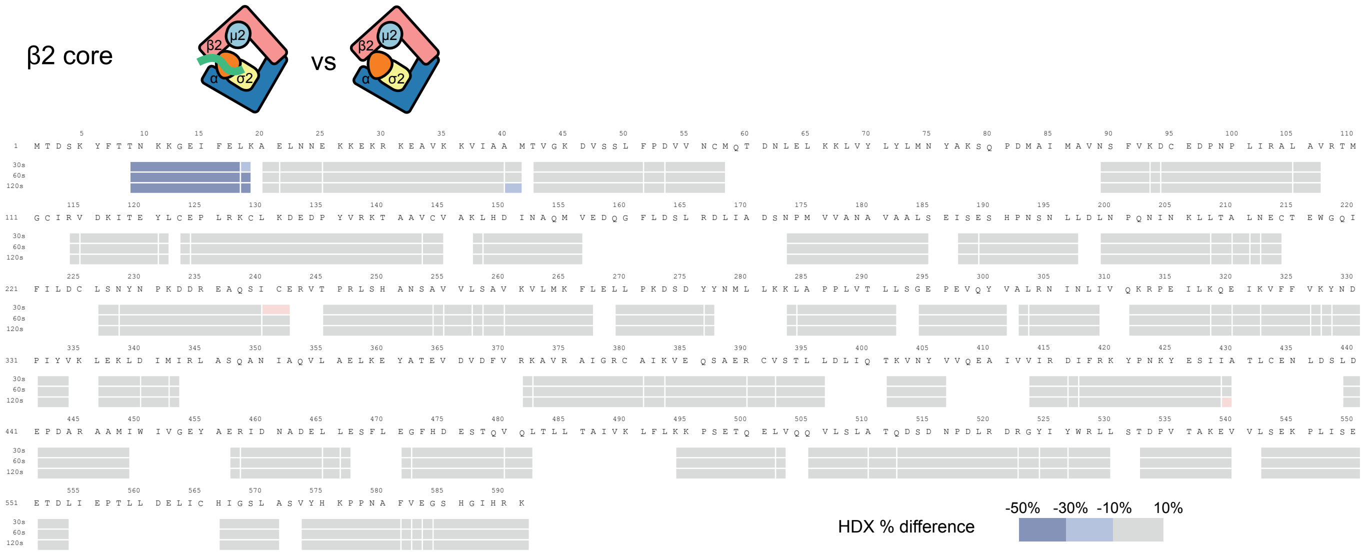

### Figure S5

A

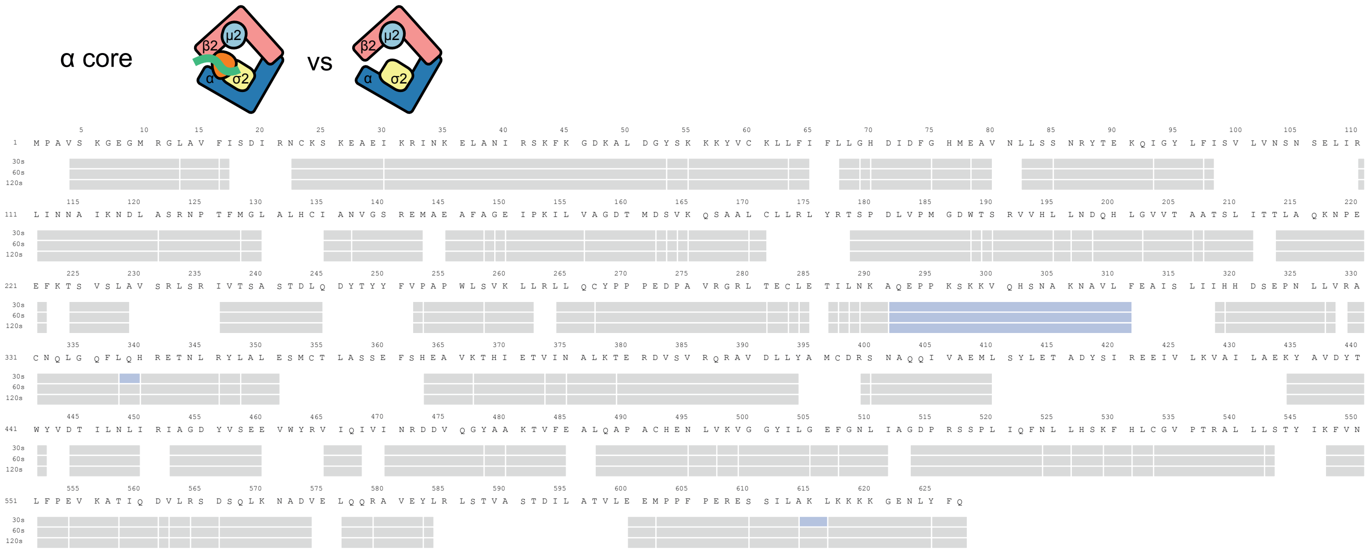
α core
β2
μ2
α
σ2
vs
β2
μ2
α
σ2

B

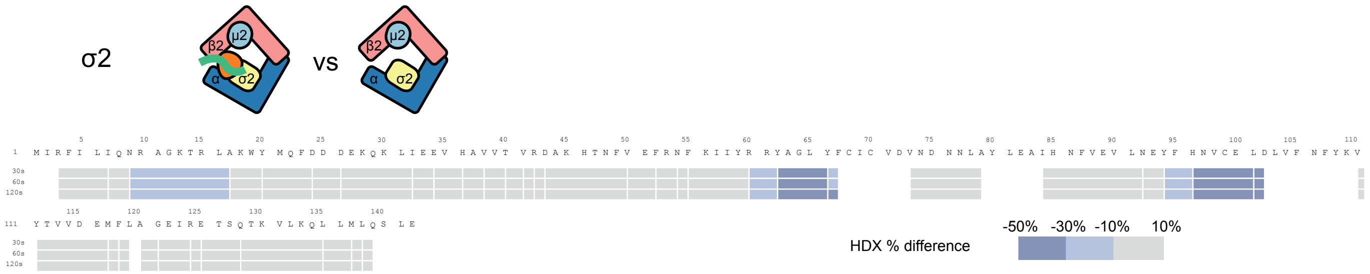
σ2
β2
μ2
α
σ2
vs
β2
μ2
α
σ2
HDX % difference
-50%
-30%
-10%
10%

### Figure S6

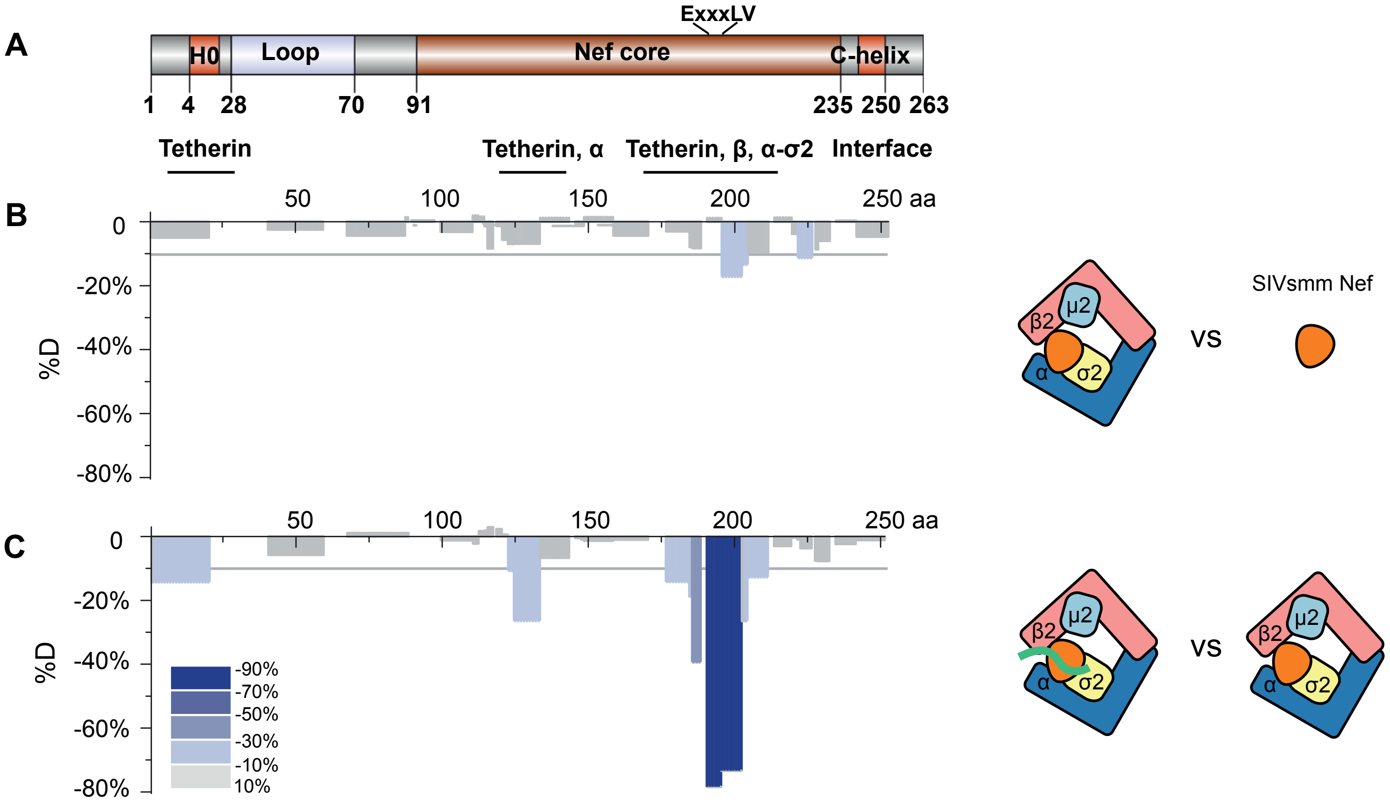
A
ExxxLV
H0
Loop
Nef core
C-helix
1
4
28
70
91
235
250
263
Tetherin
Tetherin, α
Tetherin, β, α-σ2
Interface
B
50
100
150
200
250 aa
0
-20%
-40%
-60%
-80%
%D
β2
μ2
α
σ2
VS
SIVsmm Nef
C
50
100
150
200
250 aa
0
-20%
-40%
-60%
-80%
%D
-90%
-70%
-50%
-30%
-10%
10%
β2
μ2
α
σ2
VS
β2
μ2
α
σ2

#### Figure S7

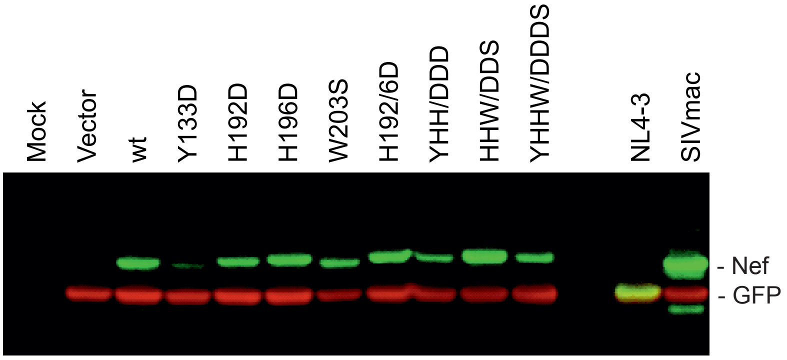
Mock
Vector
wt
Y133D
H192D
H196D
W203S
H192/6D
YHH/DDD
HHW/DDS
YHHW/DDDS
NL4-3
SIVmac
- Nef
- GFP

### Figure S8

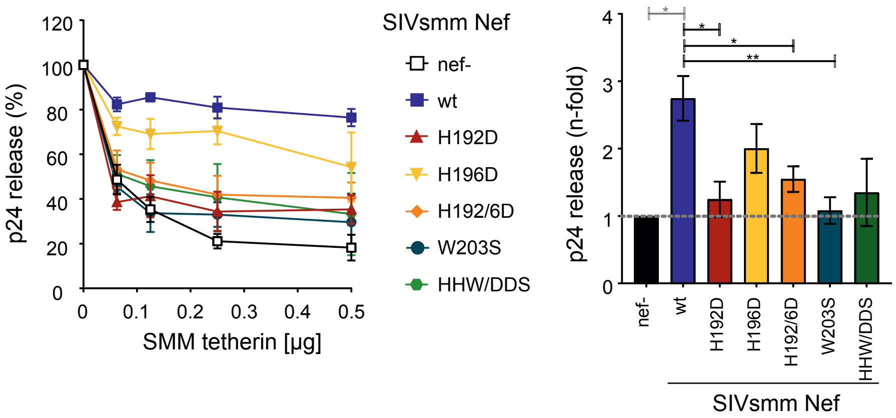

120
100
80
60
40
20
0
p24 release (%)
0
0.1
0.2
0.3
0.4
0.5
SMM tetherin [µg]
SIVsmm Nef
nef-
wt
H192D
H196D
H192/6D
W203S
HHW/DDS
4
3
2
1
0
p24 release (n-fold)
*
*
*
**
nef-
wt
H192D
H196D
H192/6D
W203S
HHW/DDS
SIVsmm Nef

### Figure S9

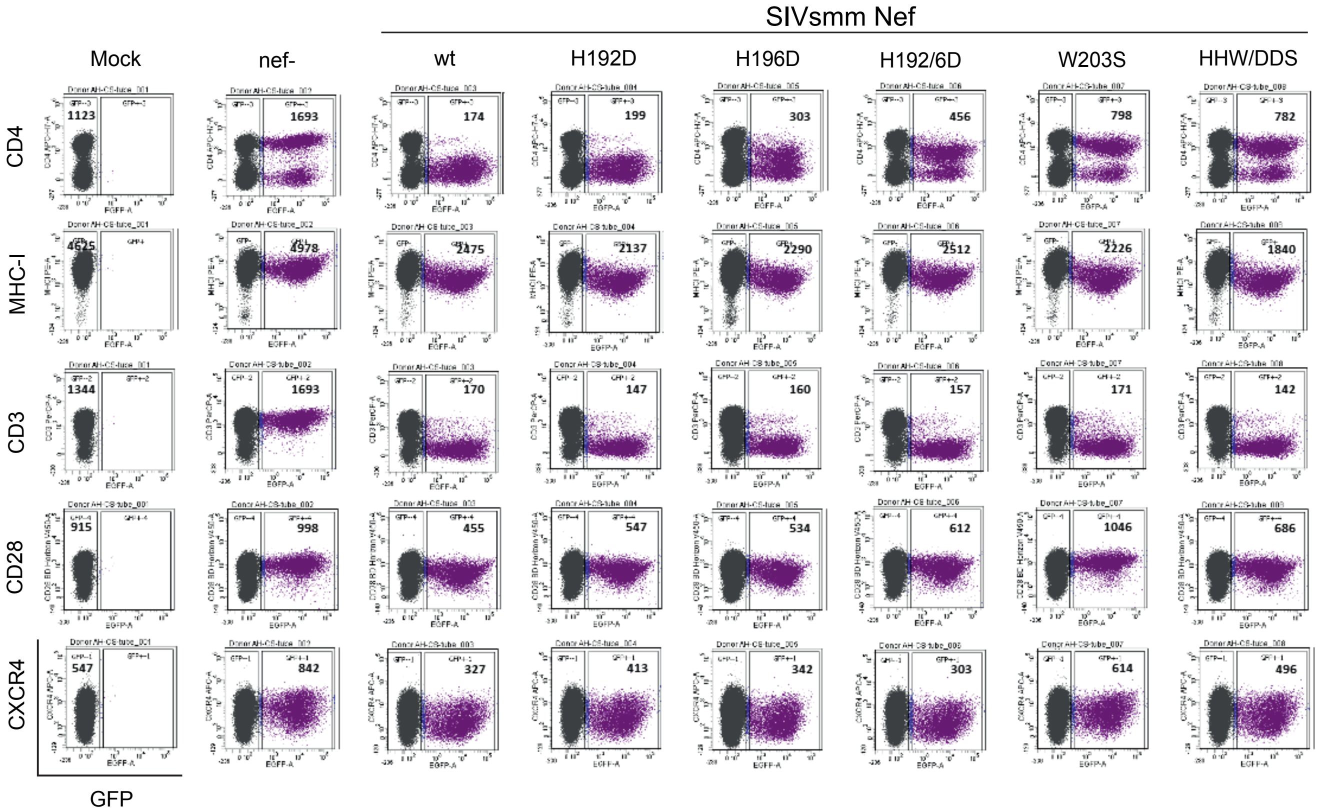
SIVsmm Nef
Mock
nef-
wt
H192D
H196D
H192/6D
W203S
HHW/DDS
CD4
MHC-I
CD3
CD28
CXCR4
GFP
1123
1693
174
199
303
456
798
782
4625
4978
2475
2137
2290
2512
2226
1840
1344
1693
170
147
160
157
171
142
915
998
455
547
534
612
1046
686
547
842
327
413
342
303
614
496
